# At least seven distinct rotavirus genotype constellations in bats with evidence of reassortment and zoonotic transmissions

**DOI:** 10.1101/2020.08.13.250464

**Authors:** Ceren Simsek, Victor Max Corman, Hermann Ulrich Everling, Alexander N. Lukashev, Andrea Rasche, Gael Darren Maganga, Tabea Binger, Daan Jansen, Leen Beller, Ward Deboutte, Florian Gloza-Rausch, Antje Seebens-Hoyer, Stoian Yordanov, Augustina Sylverken, Samuel Oppong, Yaw Adu Sarkodie, Peter Vallo, Eric M. Leroy, Mathieu Bourgarel, Kwe Claude Yinda, Marc Van Ranst, Christian Drosten, Jan Felix Drexler, Jelle Matthijnssens

## Abstract

Bats host many viruses pathogenic to humans, and increasing evidence suggests that Rotavirus A (RVA) also belongs to this list. Rotaviruses cause diarrheal disease in many mammals and birds, and their segmented genomes allow them to reassort and increase their genetic diversity. Eighteen out of 2,142 bat fecal samples (0.8%) collected from Europe, Central America and Africa were PCR-positive for RVA and 11 of those were fully characterized using viral metagenomics. Upon contrasting their genomes with publicly available data, at least 7 distinct bat RVA genotype constellations (GCs) were identified, including evidence of reassortments and 6 novel genotypes. Some of these constellations are spread across the world, whereas others appear to be geographically restricted. Our analyses also suggest that several unusual human and equine RVA strains might be of bat RVA origin, based on their phylogenetic clustering, despite varying levels of nucleotide sequence identities between them. Although SA11 is one of the most widely used reference strains for RVA research and forms the backbone of a reverse genetics system, its origin remained enigmatic. Remarkably, the majority of the genotypes of SA11-like strains were shared with Gabonese bat RVAs, suggesting a potential common origin. Overall, our findings suggest an underexplored genetic diversity of RVAs in bats, which is likely only the tip of the iceberg. Increasing contact between humans and bat wildlife will further increase the zoonosis risk, which warrants closer attention to these viruses.

**Importance:** The increased research on bat coronaviruses after SARS-CoV and MERS-CoVallowed the very rapid identification of SARS-CoV-2. This is an excellent example of the importance of knowing viruses harbored by wildlife in general and bats in particular, for global preparedness against emerging viral pathogens. The current effort to characterize bat rotavirus strains from 3 continents shed light on the vast genetic diversity of rotaviruses and also hinted at a bat origin for several atypical rotaviruses in humans and animals, implying that zoonoses of bat rotaviruses might occur more frequently than currently realized.

## INTRODUCTION

Rotaviruses are the leading cause of diarrheal disease in the young of mammals and birds. In humans, rotaviruses are responsible for 122,000-216,000 deaths in under 5-year old infants on a yearly basis, mainly in developing countries (1). The *Rotavirus* genus belongs to the family *Reoviridae* and contains 9 species designated as A-I (RVA-RVI). The rotavirus genome consists of 11 dsRNA segments encoding 6 structural viral proteins (VP1-6) and 6 non-structural proteins (NSP1-6) (2).

The RVA outer capsid antigens, VP4 and VP7 are used for a dual classification system defining P-genotype (VP4 is <u>P</u>rotease sensitive) and G-genotype (VP7 is <u>G</u>lycosylated), respectively (2). However, as gene reassortment is a common phenomenon for viruses with a segmented genome after co-infection, a more comprehensive classification approach became necessary to better account for the genome evolution and genetic diversity of RVAs. In 2008, a nucleotide sequence-based, complete genome classification system was developed for RVA, define genotypes for each of the 11 gene segment. These genotypes allowed extending the dual classification to full ‘genotype constellations’ classification (3, 4). The gene assignments are reported as Gx-P[x]-Ix-Rx-Cx-Mx-Ax-Nx-Tx-Ex-Hx, where ‘x’ denotes the particular genotype. The Rotavirus Classification Working Group (RCWG) was formed in order to assign new genotypes to rotavirus genes which could not be assigned to an established genotype (5).

Accumulating whole genome sequencing data demonstrate that there are typical GCs present in most animal species. Two of them, Wa-like and DS-1-like, are responsible for most of the human disease and designated as I1-R1-C1-M1-A1-N1-T1-E1-H1 and I2-R2-C2-M2-A2-N2-T2-E2-H2, respectively, for the non-G/P genotypes (3). Furthermore, various animal species are known to have specific GCs such as I2-R2-C2-M2-A3/A11/A13-N2-T6-E2-H3 for cattle and other even-toed ungulates (6), I1/I5-R1-M1-A1/A8-N1-T1/T7-E1-H1 for swine (3, 7), I2/I6-R2-C2-M3-A10-N2-T3-E2/E12-H7 for horses (8), and I3-R3-C3-M3-A3/A9-N2-T3-E3-H3/H6 for cats and dogs (9). Partially shared genotype patterns between established GCs, such as Wa-like human RVA strains and porcine RVAs, as well as DS-1-like human RVA strains and bovine RVAs, suggest a common origin and important zoonotic transfer events in the past (3).

Bats belong to the Chiroptera order, which is the second largest order of mammals (10). They harbor a high diversity of viruses, among them are also zoonotic viruses such as lyssavirus, Hendra and Nipah viruses, filovirus and several coronaviruses (11–17). Given their great population densities, migration ability and proximity to human habitats, bats are often screened for emerging and re-emerging viral pathogens (18, 19). Such screenings have resulted in the sporadic identification of rotavirus strains in bats in the last decade. Even though there are reports of RVH in South Korean bats (20) and Cameroonian bats (21), and a novel rotavirus species (tentatively named RVJ) was identified from Schreiber’s bats in Serbia (22); RVA is the most commonly detected species and there are currently more than 20 bat RVA strains identified. In 2010, Esona and colleagues reported the first partially sequenced RVA strain (RVA/Bat-wt/KEN/KE4852/07/2007/G25P[6]) in a Kenyan *Eidolon helvum* (straw-colored fruit bat), and the majority of retrieved gene segments were only distantly related to known mammalian RVA strains (23). During the subsequent decade, sporadic and scattered reports have been published about RVA strains in bats collected from serum, gut and fecal samples in insectivorous and fruit bats. Several of these reports came from Chinese studies (24–27), but bat RVAs were also detected and (partially) characterized from France (28), Brazil (29), Zambia (30, 31), Cameroon (32), Kenya (33) and Saudi-Arabia (34). These studies investigate samples from a variety of bat families such as Rhinolophidae (24, 30), Hipposideridae (25, 26), Vespertilionidae (26, 28), Molossidae, Phyllostomidae (29), Emballonuridae (26, 33), Pteropodidae (23, 31–34), Rhinopomatidae (34). From some of these novel bat RVA strains a few gene segments were sequenced, whereas other strains were sequenced completely, often resulting in one or multiple novel genotypes (23, 26, 29, 31, 32).

Even though RVAs are generally considered to have a rather restricted host range, a number of unusual strains have been described in literature, suggestive of interspecies transmissions involving bat RVA strains. One example is the RVA/Horse-wt/ARG/E3198/2008/G3P[3] strain that was isolated from a diarrheic foal in Argentina in 2008 (35). Although its GC was distantly related to feline/canine-like RVA strains at that time, 2 more recent publications showed a closer relationship with Chinese bat RVA strains in several gene segments (24, 25). A second example was the unusual human G3P[3] RVA strain RVA/Human-wt/JPN/12638/2014/G3P[3], isolated from a 4 year-old child with severe gastroenteric symptoms in Japan. Three out of its 11 gene segments were closely related to a South African bat RVA strain, suggesting a reassortment involving a bat RVA strain (36). A third example are two unique G20 human RVA strains, RVA/Human-wt/ECU/Ecu534/2006/G20P[28] (37) and RVA/Human-wt/SUR/2014735512/2013/G20P[28] (38). The recent identification of the G20 genotype in a Brazilian bat RVA strain (RVA/Bat-wt/BRA/3081/2013/G20P[x]) also suggests a potential bat reservoir for these human strains (29).

All in all, slowly emerging data on bat RVA strains start to show that some unusual human and animal RVA strains might actually have been derived from bats. Therefore, the global surveillance of novel and reassortant RVA bat strains has to continue in order to better understand the genetic diversity of bat RVA strains, as well as to maintain both public and animal health. Here we report identification of 11 bat RVA strains from Bulgaria, Gabon, Ghana and Costa Rica, suggesting evidence of multiple reassortment and host switching events from bats to bats and to other mammals.

## RESULTS & DISCUSSION

Bats are known hosts of various human pathogens, including viruses such as rabies virus, henipaviruses, Marburg virus, SARS and MERS CoVs (11–17). In addition, there have been sporadic reports on several other RNA viruses in bats such as paramyxoviruses, picornaviruses, orthoreoviruses and astroviruses (39–42). Bat rotaviruses have also been sporadically reported during the last decade and it was rotavirus A (RVA) that has been the most frequently reported rotavirus species. This is not very surprising given the fact that RVA has been detected in a wide range of mammals and birds (43–45). Furthermore, there are plenty of examples of this enteric pathogen being capable of interspecies transmission in literature, sometimes in combination with reassortments, between various mammalian species including humans (46). In some occasions, such animal-derived gene segments (e.g. VP7 genotypes G8 from cattle, G9 and presumably G12 from pigs) or complete GCs (AU-1 like strains from cats) have become established in the human population. This established circulation either happened in a limited geographical region (AU-1 like or G8) or worldwide; such as epidemiologically important human pathogenic G9 and G12 RVAs (47, 48).

In order to further investigate the potential of bat RVA strains to spill over between bats or towards other mammalian species, we investigated RVA strains from over 2,000 bats, spanning 5 countries in 3 continents. The bat fecal samples that were collected from Bulgaria, Romania, Germany, Gabon, Ghana and Costa Rica were screened for RVA, using a nested RT-PCR targeting a short piece of the highly conserved polymerase gene (VP1, Table S1). This screening yielded 18 positives out of the 2,142 screened samples (0.8%) (Table S2). The RVA detection rate per species ranged from 0 to 1.1%, except for *H. gigas* (14.9%). The reason for this higher detection rate is unknown, but could be due to: 1) better matching oligonucleotides used for the detection, 2) an ongoing RVA outbreak in the sampled caves, or 3) higher circulation of enteropathogens in *H. gigas*. RVA positive samples were collected from five bat families Pteropodidae, Rhinolophidae, Hipposideridae, Phyllostomidae and Vespertilionidae, and they originated from all sampling sites except Romania.

### Eleven near complete bat RVA genomes, including 6 novel genotypes

From 16 of the RVA positive samples, a sufficient amount of sample was available for complete viral genome sequencing using the NetoVIR protocol (Table S3). 118.9 million paired-end (PE) reads (2×150 base pairs) and an average of 7 million PE reads/sample were generated by Illumina sequencing (Table 1). Four samples from Gabon and 1 sample from Germany did not yield any RVA contigs longer than 500 base pairs and were therefore not investigated further. From 11 samples near complete RVA genomes could be retrieved. These RVA samples belonged to 5 out of the 46 tested bat species (10.8%), from 4 out of the 10 (40%) tested families, as shown in the bat phylogenetic tree (Table S2, Figure S1). The percentage of reads mapping to RVA in each sample ranged from 0-90% (Table 1).

**Table 1.**
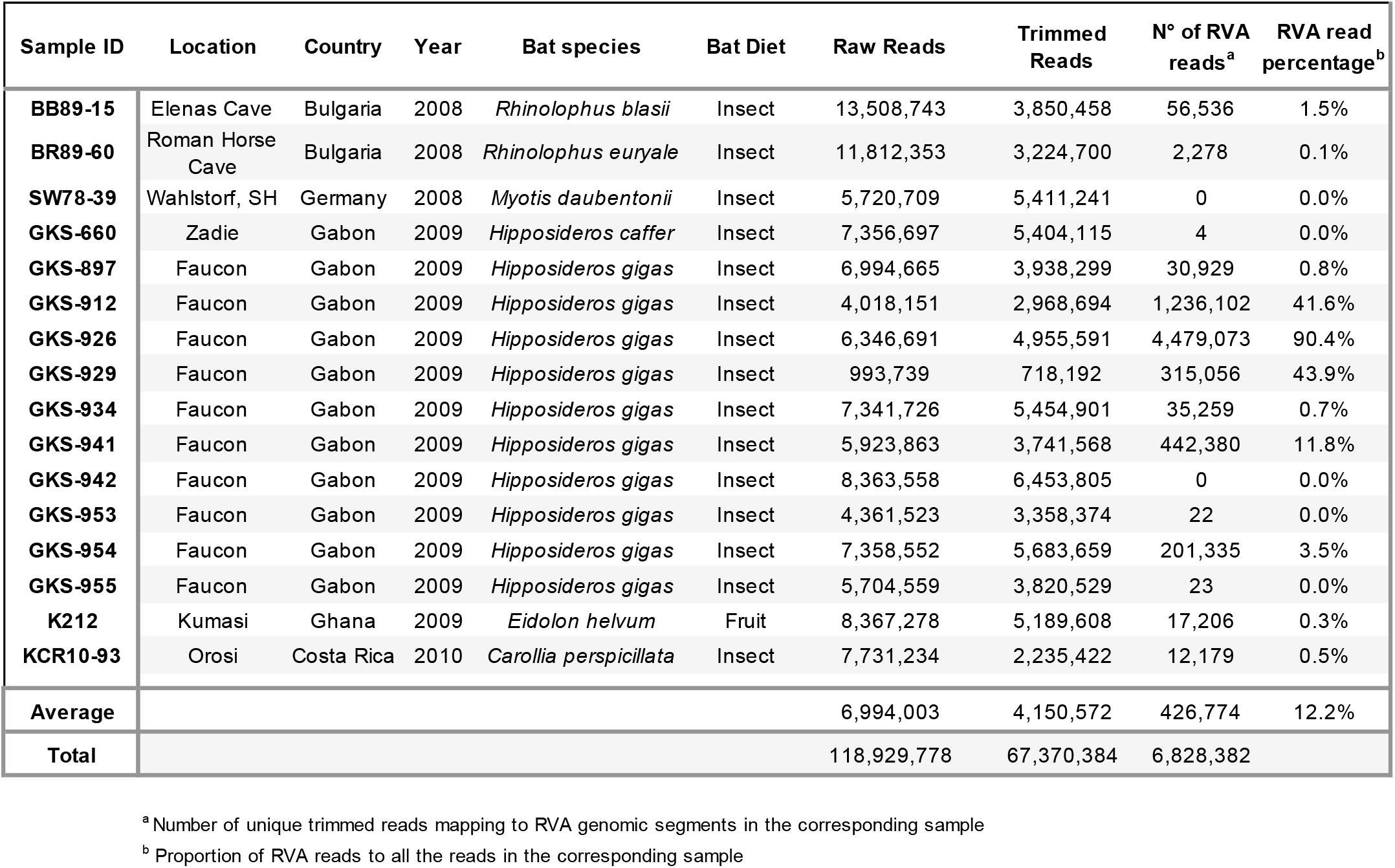
Meta-data and NGS summary of the sequenced RVA-positive samples

The GCs of the 11 bat RVA strains are shown in Table 2. The genotype assignments, including novel VP6 (I30), NSP1 (A36), NSP2 (N23) and NSP4 (E28) genotypes for some of the Gabonese strains and NSP1 (A32) and NSP3 (T23) genotypes for the strain from Costa Rica were made according to the guidelines determined by the RCWG (49). Although the NSP5 gene segment of RVA/Bat-wt/CRC/KCR10-93/2010/G20P[47] most likely also represents a novel genotype, we were not able to retrieve the complete ORF (despite several attempts using RT-PCR and Sanger sequencing), which is required for the assignment of a novel genotype (50). Particular GCs were identified in different geographic locations (Table 2). Gabonese strains were similar to each other, with certain genotypes shared with the Bulgarian strains (G3, P[3], C3, M3, N3, T3 and E3). However, they do not cluster phylogenetically closely together (*vide supra*), indicating non-recent reassortment events. KCR10-93 also possessed a unique GC, except for the VP4 genotype P[47], which was shared with the Ghanaian strain. Interestingly, these 2 VP4 genes were very closely related (*vide supra*), suggesting a recent reassortment event. Gabonese GKS-912, GKS-926 and GKS-934 appeared to have a co-infection, as multiple genotypes were identified in these samples for VP2, VP3, VP4, NSP2, NSP3 and NSP4. For GKS-934, 2 near complete VP7 gene segments were identified, both belonging to the G3 genotype, yet having a substantial nucleotide level dissimilarity (19%, *vide infra*). This was also the case for K212 possessing 2 distinct M14 genotypes with 12% nucleotide sequence distance.

**Table 2.**
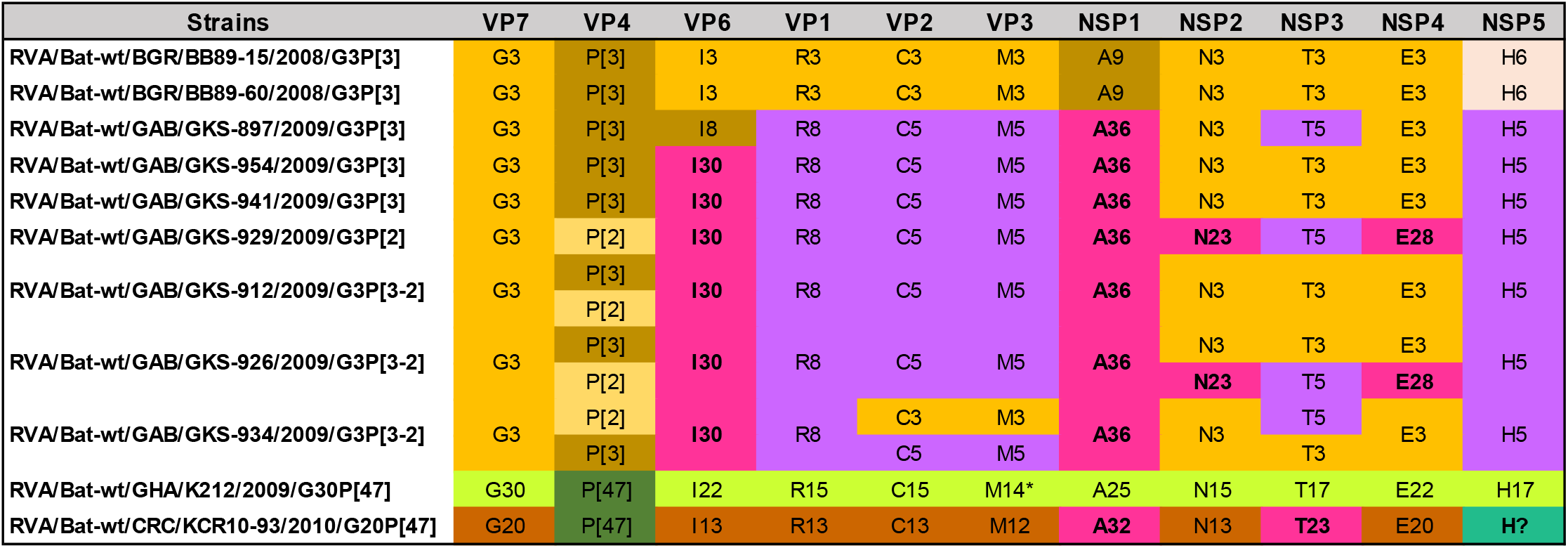
Color-coded GCs of the bat RVA strains identified in this study. In some samples, 2 different variants of the same gene segments were identified, suggesting co-infections. K212 possessed 2 distinct VP3 gene segments belonging to the same M14 genotype (indicated with an asterisk). NSP5 gene of KCR10-93 could not be assigned to any of the established genotypes; neither assigned to a novel genotype as the complete ORF could not be determined. Therefore, this genotype is indicated as “H?”.

### At least 7 seven distinct bat RVA genotype constellations

Even though most animal species, including humans, have a limited number of typical RVA GCs, the RVAs harbored by bats show a great genetic diversity. Combining our data with previously published bat RVA genomes showed that there are at least 7 distinct bat RVA GCs circulating in the bat population (Table 3), ranging from completely unique to partially overlapping with each other. The Bulgarian RVA/Bat-wt/BGR/BB89-15/2008/G3P[3] and RVA/Bat-wt/BGR/BR89-60/2008/G3P[3] strains were identical or very similar to MSLH14-like RVA strains from China and a partially sequenced strain from Brazil (“orange” GC in Table 3). Even though at least 3 of the samples from Gabon possessed more than one RVA strain, they possessed at least 3 distinct but related GCs (“purple” GC in Table 3), not previously identified in bats. RVA/Bat-wt/GHA/K212/2009/G30P[47] (“green” GC in Table 3) was identical or very similar to several previously identified Cameroonian bat RVA strains (32), as well as some partially sequenced bat RVA strains from Zambia (31). RVA/Bat-wt/CRC/KCR10-93/2010/G20P[47] had a distinct GC (“brown” GC in Table 3), including at least 2 previously undescribed genotypes, and shared the G20 genotype with RVA/Bat-wt/BRA/3081/2013/G20P[x]. Of interest was the P[47] genotype, which was shared with 2 African strains from the green GC. The “yellow” GC in Table 3 was composed of 2 strains with identical genotypes; RVA/Bat-wt/CMR/BatLy03/2014/G25P[43] and RVA/Bat-wt/SAU/KSA402/2012/G25P[43], detected in Cameroon and Saudi Arabia, respectively, as well as the partially sequenced strain RVA/Bat-wt/KEN/KE4852/2007/G25P[6] from Kenya. Two GCs (indicated in “blue” and “dark grey” in Table 3) were only represented by a single bat strain from Kenya (RVA/Bat-wt/KEN/BATp39/2015/G36P[51]) and China (RVA/Bat-wt/CHN/GLRL1/2005/G33P[48]), respectively (Table 3).

**Table 3.**
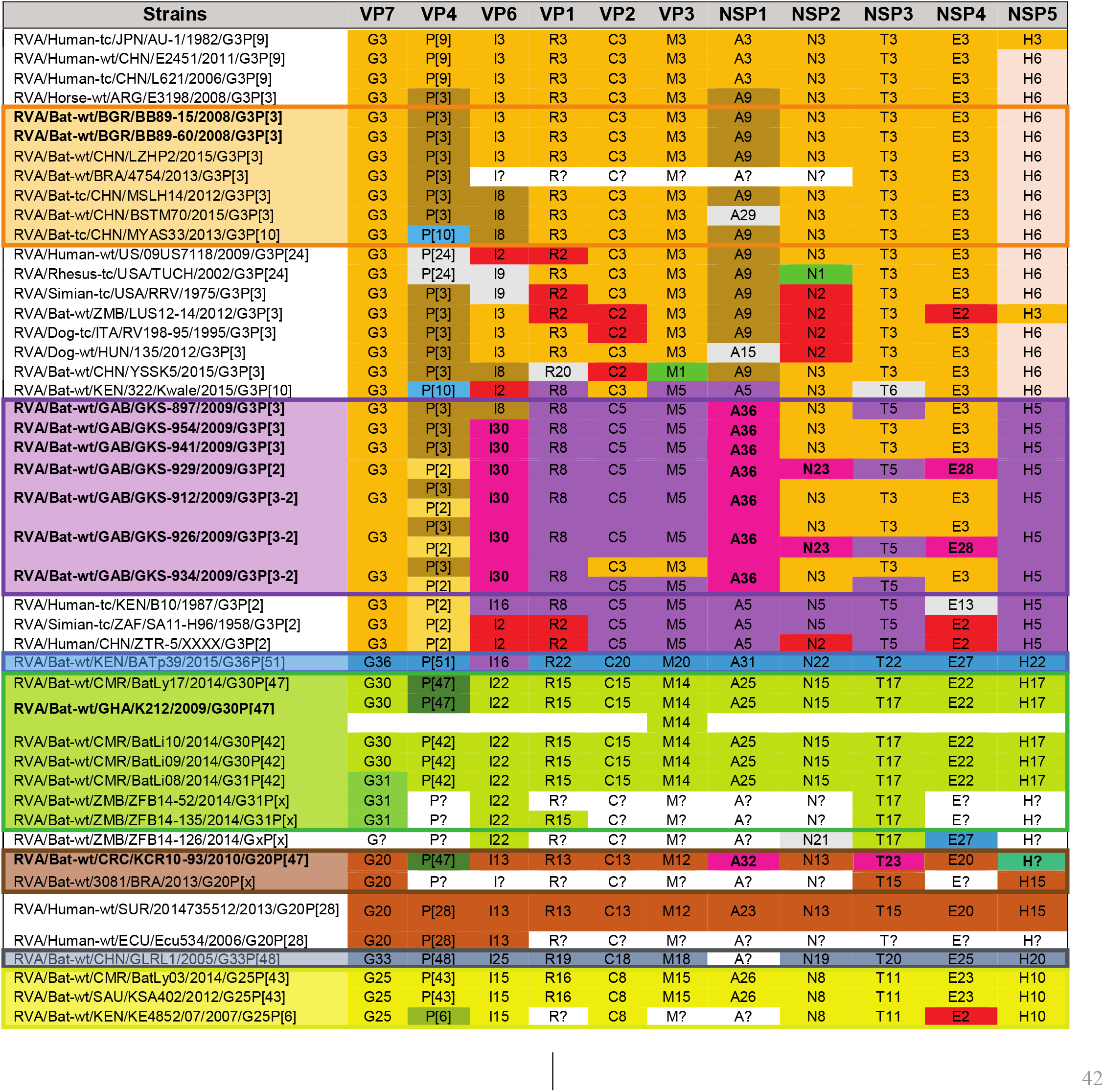
Color-coded GCs for the bat RVA strains identified in this study, previously published bat RVA strains, as well as a selection of RVA strains from other host species potentially related to bats. The non-sequenced segments or unassigned genotypes are denoted with ‘[letter code]?’. The genotypes colored in light grey are less relevant due to a lack of (in)direct genomic relationship with bat RVAs identified in the current study. The strain names are color-matched with the corresponding GCs (orange, purple, blue, green, brown, dark grey and yellow).

### Reassortments among bat RVA strains

Even though the GCs are somewhat conserved, there are ample examples for the occurrence of reassortments. In the orange GC, there are some unusual genotypes such as P[10] for VP4, R20 for VP1 and A29 for NSP1 (Table S4a and S4b), which are most likely the results of reassortment events with currently unknown RVA strains (25, 26). Reassortment also takes place between different bat RVA GCs, albeit to a limited extension. For example, RVA/Bat-wt/GAB/GKS-897/2009/G3P[3] is the only strain from the purple GC with the I8 VP6 genotype, which is shared with several strains from the orange GC (RVA/Bat-tc/CHN/MSLH14/2012/G3P[3], RVA/Bat-wt/CHN/BSTM70/2015/G3P[3], RVA/Bat-tc/CHN/MYAS33/2013/G3P[10] and RVA/Bat-wt/CHN/YSSK5/2015/G3P[3]), suggesting a reassortment event. A second example is the shared P[47] VP4 genotype between RVA/Bat-wt/GHA/K212/2009/G30P[47] and RVA/Bat-wt/CMR/BatLy17/2014/G30P[47] (green GC) and RVA/Bat-wt/CRC/KCR10-93/2010/G20P[47] (brown GC) (Table 2). Interestingly, these last 3 strains were 97-100% identical to each other on the nucleotide level for VP4, suggesting a recent reassortment event. Finally, there are also a few bat RVA strains with unusual genotype composition, which do not clearly fall into the 7 described GCs. RVA strains RVA/Bat-wt/ZMB/LUS12-14/2012/G3P[3] and RVA/Bat-wt/CHN/YSSK5/2015/G3P[3] possess several genotypes typical for the orange GC, in addition to several other genotypes of unknown origin (Table S4b). Finally, RVA/Bat-wt/KEN/322/Kwale/2015/G3P[10] possesses both genotypes typical to the orange and purple GCs, in addition to some atypical bat RVA genotypes.

### RVA interspecies transmission in bats and potential host range restriction

As demonstrated by the orange GC, RVAs belonging to certain bat families might undergo multiple host switching events. The Bulgarian RVA strains were isolated from rhinolophid bats, whereas the Chinese MSLH14-like strains were found in bats from the Rhinolophidae, Hipposideridae and Emballonuridae families (Table S4a).

In addition to RVAs potentially being able to infect multiple bat families, individual bat families could also harbor more than one GC, as is shown in Table S4c. Pteropodid bats harbor completely unique GCs (green and yellow), suggesting that the associated RVA strains have a high epidemiologic fitness in these populations. This further indicates that the Pteropodidae, which includes the straw-colored fruit bats, has been a substantial virus reservoir for a long time already, as also shown for Marburg virus, Hendra and Nipah viruses (12–14).

### Wide geographic dispersal of bat RVA GCs

The global distribution of the bat RVA GCs revealed several patterns regarding RVA circulation in bats, as shown in Figure 1. Bat RVAs belonging to the brown, purple, blue and dark grey GCs have so far only been identified in Costa Rica (and perhaps Brazil), Gabon, Kenya and China, respectively. On the other hand, the green and yellow GCs were confirmed to be further dispersed, from Cameroon to Saudi Arabia (G25P[43]), and from Ghana and Cameroon to Zambia, respectively, as was previously suggested by Sasaki *et al.* (51). However, highly similar RVA strains belonging to the orange MSLH14-like GCs span at least 3 different continents and subcontinents, e.g. Asia, Europe and possibly Central America. Furthermore, it was also shown that RVA strains with distinct GCs could co-circulate in the same region, as is the case in Cameroon (green, yellow and purple GCs) and China (orange and dark grey GCs) (Figure 1).

**Figure 1.**
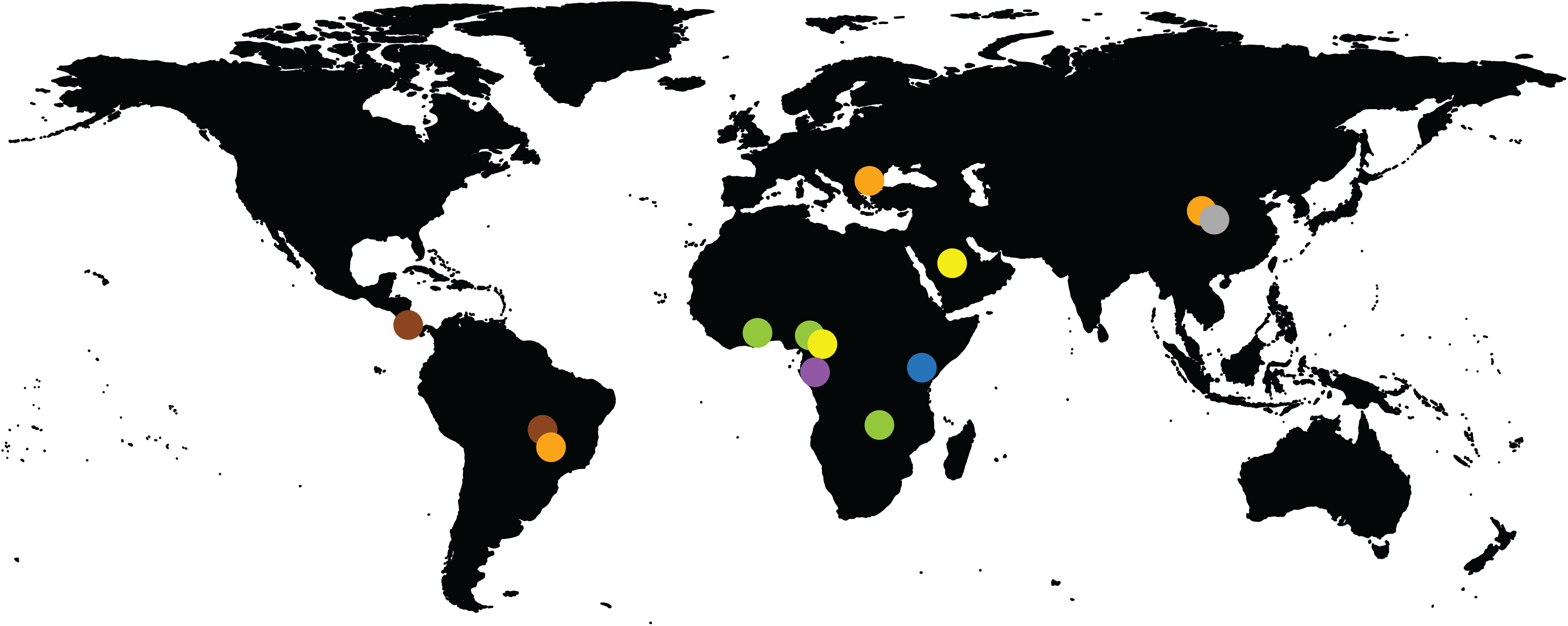
Geographic distribution of the currently known bat RVA GCs. The colored dots on the map represent the circulating genotypes at the specified locations according to the GCs shown in Table 3.

With powered flight, migratory bats can travel long distances between summer and winter roosts, for foraging and searching for a mate (52). Among long-distance migratory bats, *E. helvum* can cover a range of 270 to 2,500 km (53), vespertilionid ‘tree bats’ and the subtropical/tropical molossid bats can fly over 1,000 km (54, 55). Global distribution and intercontinental bat virus transfers are also typical to other bat viruses (56). In addition to migration across vast distances, the fact that some distinct GCs seem to have overlapping geographical ranges (such as in China and West Africa in Figure 1) suggest a fitness advantage for these particular genotypes occurring together. However, there is also ample evidence of gene reassortment events among established GCs (e.g. P[47] in green and brown GC; or I8 in purple and orange GC), or with RVA strains of currently unknown origin (e.g. A29, A15, E27).

It is clear that more bats should be sampled in order to have a comprehensive understanding of the driving and restricting forces of bat RVA genetic diversity, or the lack thereof. The detection of P[47] reassortment between Ghanaian and Costa Rican bat RVAs, which are located more than 9,000 km’s apart, cannot only be explained by the flight ability of bats, but rather the lack of sampling between these 2 locations. We hypothesize that with the increasing bat RVA sequencing efforts, the geographical and host range of most GCs (such as the blue, dark grey, yellow and brown) will be significantly expanded.

### Potential of interspecies transmissions of bat RVA to mammalian hosts

We further investigated whether there is potential for unusual RVA strains detected in other mammals (including humans) to be a result of an interspecies transmission from bat strains identified in the current and other studies (Table 3).

#### Likely transmission of bat RVA strains to a horse

In 2013, Miño and colleagues reported an unusual Argentinian equine G3P[3] RVA strain RVA/Horse-wt/ARG/E3198/2008/G3P[3]. Based on the GC, it was speculated to have a common ancestor with both feline/canine RVA strains, as well as the unusual rhesus RVA strain RRV. However, the nucleotide identities were below the 90% for most of the genome segments, suggesting that the original host may not be identified yet (35). When more bat RVA genomes became available in subsequent years, Xia and colleagues, and later also Biao He and colleagues, suggested that E3198 might be of bat origin, based on the GCs and nucleotide similarities (25, 26). The close genetic relationship between E3198 and the Bulgarian strains presented here, across all 11 gene segments, might further suggest a bat origin of this unusual equine RVA strain (Figures 2–4, Figure S2a, nucleotide similarities 87-97%).

**Figure 2.**
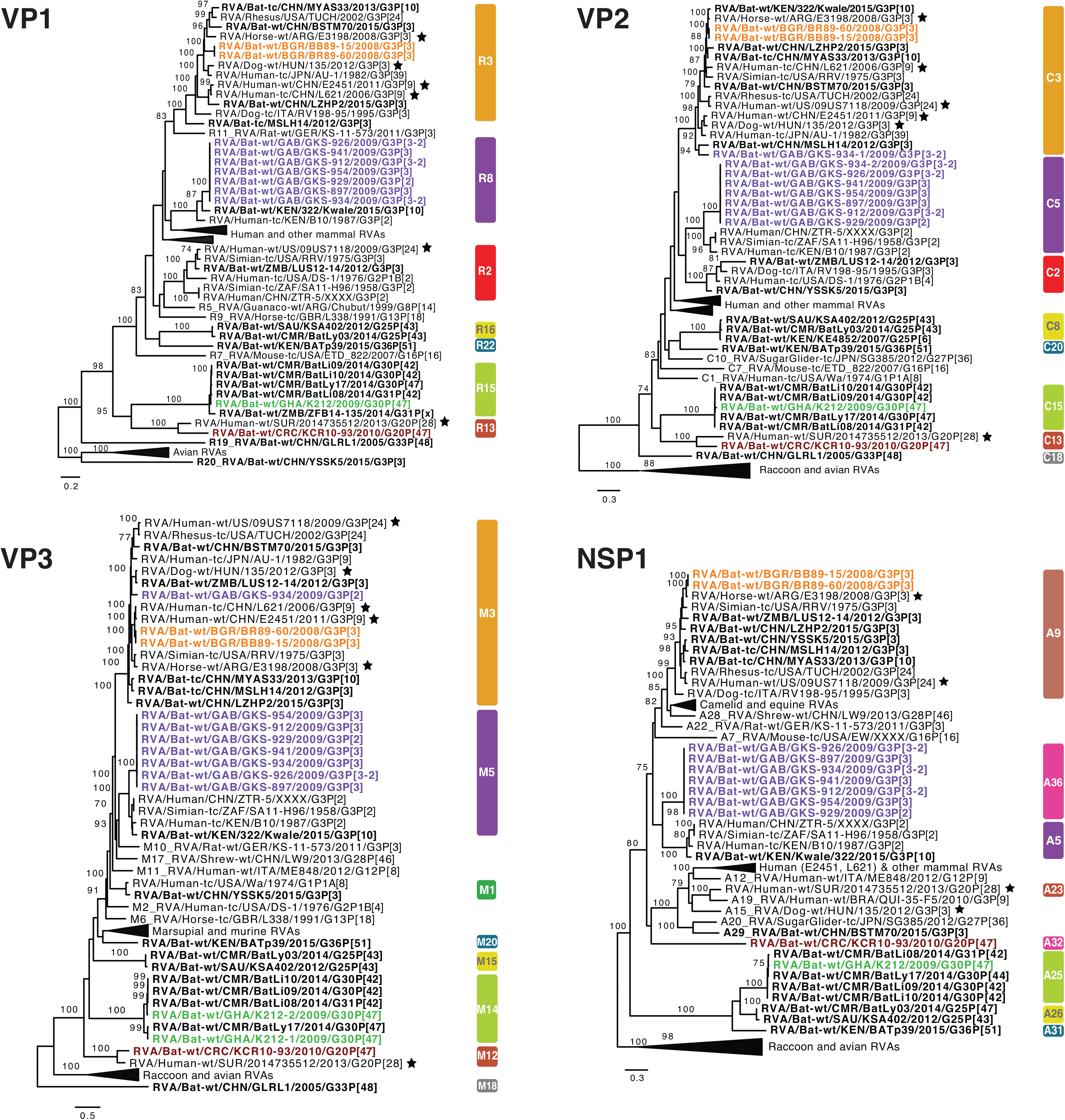
Maximum likelihood trees of the VP1, VP2, VP3 and NSP1 genes of the identified bat RVA strains with known human, bat and other mammal RVAs. Only bootstrap values above 70 are shown. The genotypes are listed on the right side of the trees. The bat RVA strains identified in this study are shown in bold and colored to their GC, previously reported bat RVA strains are shown in bold in black, and non-bat RVA strains related to a bat RVA strain are marked with filled stars.

#### Unexpected high similarities between bat and simian RVA strains

RVA strain RVA/Simian-tc/ZAF/SA11-H96/1958/G3P[2] was isolated from an overtly healthy vervet monkey and has subsequently been used extensively as a laboratory strain in RVA growth, virulence, genome replication and in recent years also a reverse genetics research (57–59). However, its origin remained obscure, as related strains were never identified in vervet monkeys or other non-human primates ever after. In 2011, Ghosh and colleagues identified an unusual RVA strain RVA/Human-tc/KEN/B10/1987/G3P[2] from a child in Kenya, which shared 8 out of 11 genotypes with SA11-H96. They speculated about a simian or other animal origin of this strain (60). Around the same time, a second human RVA strain RVA/Human/CHN/ZTR-5/XXXX/G3P[2], nearly identical to SA11-H96 (Figure S2b) was deposited in GenBank as a potential vaccine candidate. However, the controversy about the origin of these SA11-like strains (SA11-H96, B10 and ZTR-5) remained. To our surprise, the purple GC described in this paper, containing only the bat RVA strains from Gabon, showed up to seven genotypes in common with these SA11-like strains (Table 3), with varying degrees of nucleotide similarities (Figures S2b, c). According to phylogenetic analyses the bat RVAs from Gabon and Kenya clustered with B10 for the VP1, VP6, NSP4 gene segments, and with all 3 strains (B10, SA11-H96 and ZTR-5) for VP2-4, NSP1, NSP3 and NSP5 (Figures 2–4).

Not only for SA11-H96, but also for RVA/Simian-tc/USA/RRV/1975/G3P[3] and RVA/Rhesus-tc/USA/TUCH/2002/G3P[24], some close relationships with bat RVA strains were noted. The VP1, VP3, VP4, VP6, VP7, NSP1-5 genes of RRV clustered closely with one or multiple bat and bat-related RVA strains (Figures 2–4). For TUCH, the VP1, NSP1, NSP5 gene segments also clustered close to bat RVA strains (Figures 2–4).

The finding that the purple SA11-like GC was found in multiple bats in Gabon, and only on a single occasion in vervet monkeys and in 2 unrelated human cases, makes bats the prime suspect of being the major hosts of these viruses, making the monkey and humans strains putative examples of interspecies transmissions. It should however be noted that the phylogenetic clustering between these bat, simian and human strains is still rather variable, and the nucleotide similarities are not as high as between bat RVA strains and RVA/Horse-wt/ARG/E3198/2008/G3P[3] (Figure S2a, Figures 2–4), suggesting that more RVAs from currently unsampled animal species will likely cluster in between. However, 2 other bat strains are of further interest: 1) the bat RVA strain RVA/Bat/KEN/322/Kwale/2015/G3P[10] (only available as a GenBank entry at this point) seems to have a mixed GC possessing both characteristics of the orange and purple GCs (Table 3). Especially, the purple genotypes R8, M5 and A5 of 322/Kwale are of interest as they are much more closely related to the SA11-like strains than the Gabon bat RVA strains (Figure 2); 2) the bat RVA strain RVA/Bat-wt/KEN/BATp39/2015/G36P[51] (only available in GenBank) possesses a single purple genotype I16, and again this is more closely related to the SA11-like strain B10 compared to the Gabon RVA strains. Taken all together, we speculate that with further RVA screenings in bat populations, more bat RVA strains that are closely related to the vervet monkey RVA strain SA11-H96 and human SA11-like RVA strains may be detected.

#### Evidence of bat RVA strains transmitted to humans?

The G3 genotype is usually associated with P[8] genotype in humans RVAs, and combinations such as G3P[3] and G3P[9] are only sporadically found in the human population (61). Nonetheless, in the 2000-2001 season, the VP4, VP7, VP6 and NSP4 genes were sequenced from a rare human strain RVA/Human-wt/THA/CMH222/2001/G3P[3], detected in a 2 year-old severely diarrheic patient in Thailand (41). It was reported to have a VP7 gene closely related to RVA/Simian-tc/USA/RRV/1975/G3P[3] and a VP4 gene that was caprine-like. Subsequently, Xia and colleagues speculated that this strain is distinct from typical human RVA GCs and very likely shared a common ancestor with Asian bat RVAs (33). Our study provides further evidence for the bat origin of CMH222, as the VP6 I8 genotype of CMH222 is closely related to RVA/Bat-wt/GAB/GKS-897/2009/G3P[3] (Figure 3).

**Figure 3.**
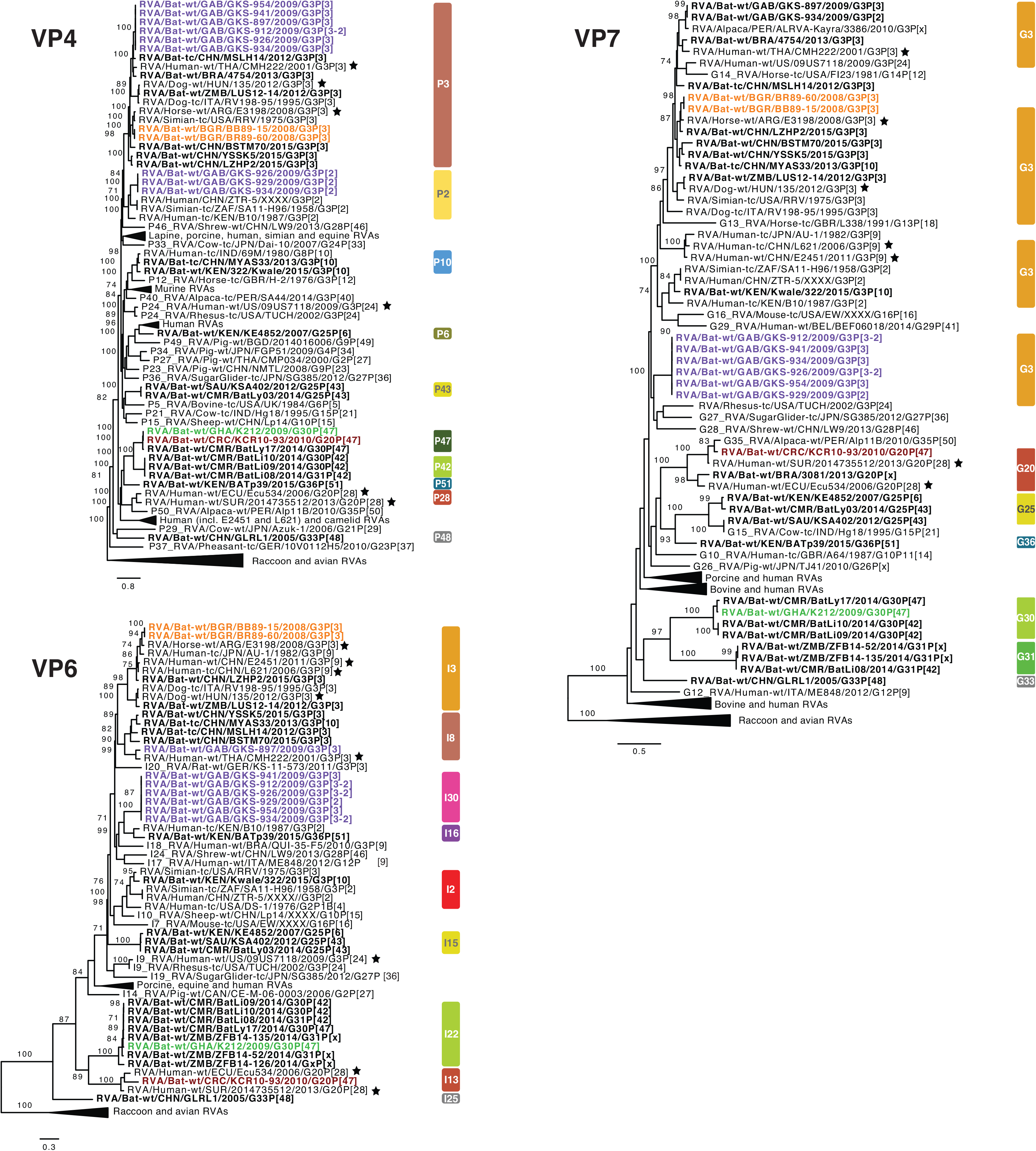
Maximum likelihood trees of the VP4, VP6 and VP7 genes of the identified bat RVA strains with known human, bat and other mammal RVAs. Only bootstrap values above 70 are shown. The genotypes are listed on the right side of the trees. The bat RVA strains identified in this study are shown in bold and colored to their GC, previously reported bat RVA strains are shown in bold in black, and non-bat RVA strains related to a bat RVA strain are marked with filled stars.

Later on, Wang and colleagues contributed to the list of unusual Southeast Asian human RVA strains. Possessing the G3P[9] genotypes, both the RVA/Human-tc/CHN/L621/2006/G3P[9] and RVA/Human-wt/CHN/E2451/2011/G3P[9] strains were isolated from a symptomatic adult and a symptomatic child, respectively (62). Complete genome analyses revealed a high genetic relatedness to strains of feline/canine origin for almost all 11 genes. L621 and E2451 also clustered near the aforementioned unusual RVA/Horse-wt/ARG/E3198/2008/G3P[3] for the VP3, VP6, NSP2, NSP5 genes; and L621 additionally also clustered with the E3198 NSP3 gene. Here, we observed that these atypical Asian human strains were also closely related to the Bulgarian bat RVA strains for VP3, VP6, NSP2, NSP4, NSP5 and Gabonese bat strains for NSP2, NSP3, NSP4 of the orange GC (Figures 2–4). These additional findings further substantiate, as well as complicate the identification of the likely bat host, from which the L621 and E2451 strains likely jumped to humans.

**Figure 4.**
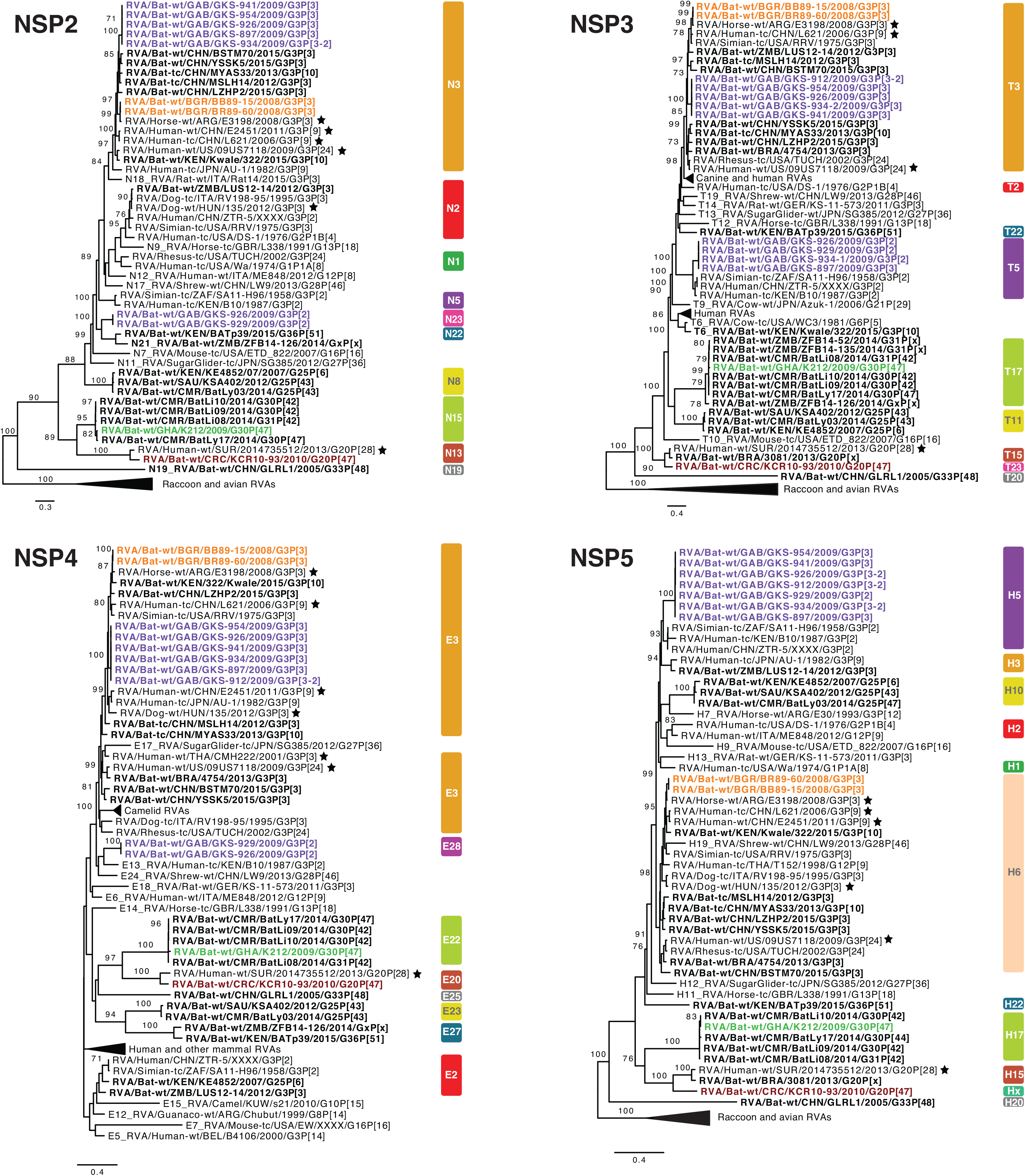
Maximum likelihood trees of the NSP2, NSP3, NSP4 and NSP5 genes of the identified bat RVA strains with known human, bat and other mammal RVAs. Only bootstrap values above 70 are shown. The genotypes are listed on the right side of the trees. The bat RVA strains identified in this study are shown in bold and colored to their GC, previously reported bat RVA strains are shown in bold in black, and non-bat RVA strains related to a bat RVA strain are marked with filled stars.

Following these potential zoonosis reports, Esona and colleagues also revealed remarkable findings in Latin America in 2018, where only limited bat RVA information is present to date (38). A human strain RVA/Human-wt/SUR/2014735512/2013/G20P[28] was isolated in Suriname, and possessed a rare G20 genotype, which was also detected in an Ecuadorian human RVA strain (Ecu534) in 2006. Remarkably, 2014735512 showed high similarities with bat strain RVA/Bat-wt/BRA/3081/2013/G20P[x] for the VP7, NSP3 and NSP5 genes (Figure S2d) and it was speculated to be of bat origin as these genotypes have not been detected in any other animal species so far. RVA/Bat-wt/CRC/KCR10-93/2010/G20P[47] also showed nucleotide similarities ranging from 82% to 92% with 2014735512 for 9 out of 11 gene segments, and also phylogenetically clustered together, albeit not very closely (Figures 2–4). Even though more evidence is needed, this finding might indicate a bat RVA origin for this rare human RVA strain.

## Conclusion

Despite the limited number of bat species that have been screened for rotaviruses, a surprisingly large genetic diversity of RVA strains is presented in this study, including 6 novel genotypes. With increasing screening efforts, it is without a doubt that this diversity will expand both genetically and geographically. We also presented multiple examples of close genetic relatedness of several mammalian and bat rotaviruses. The indicated zoonoses has - to the best of our knowledge - always been restricted to sporadic cases so far and has never resulted in major outbreaks in humans. However, it is believed that the rotavirus genotype constellations currently circulating in humans also have a common ancestor with animal rotaviruses, highlighting that interspecies transmissions followed establishment in the human population could happen again (3).

Another notable finding is that several gene segments of bat RVA strains and the simian SA11 RVA strain (the latter being used in global rotavirus research for decades), have a common origin. Furthermore, SA11 strain has been recently used as the backbone of a RVA reverse genetics system, and is therefore likely to be used even more in the future. It would be intriguing to test whether or not SA11 grows well in bat cell lines, or in *in vivo* infection experiments.

## MATERIALS AND METHODS

### Sample collection

Fecal samples were collected from 2,142 bats from 10 bat families, representing 46 bat species (Table S2). Sample collection took place in Ghana, Gabon, Bulgaria, Romania, Germany and Costa Rica during 2008-2010 as part of investigations of other viruses in bats, such as coronavirus, astrovirus, and picornavirus, as described previously (56, 63–66). Bat species were determined by trained field biologists. For European and Costa Rican studies, bats were caught with mist nets, put into cotton bags and fecal pellets are collected. Ghanaian fecal droppings were collected with plastic foil from the trees in which *E. helvum* bats were roosting. The pellets were kept in RNAlater RNA stabilization solution (QIAGEN, Hilden, Germany). Gabonese bats were also captured with mist nets just before twilight and were individually euthanized. Bat feces were collected with the corresponding permissions of the host countries in all of the studies.

### RT-PCR rotavirus screening and viral metagenomics

Viral RNA was isolated from the fecal specimens as described previously (65). To screen the RVA presence in bats, conserved RVA-specific primer pairs targeting the VP1 gene were used (277 nucleotide long PCR product) in a hemi-nested and single round reverse transcription (RT-PCR) assay (Table S1). Among the 18 positive specimens (Tables S2–S3), 16 fecal samples, of which sufficient material was left, were shipped to the Laboratory of Clinical and Epidemiological Virology, Leuven, Belgium on dry ice, for further complete genome analyses (Table 1).

The NetoVIR protocol was used for viral enrichment of the fecal suspensions as described before (67). Briefly, the fecal samples were suspended in dPBS and homogenized with a MINILYS homogenizer (Bertin Technologies) for 20s at 3,000 rpm. The homogenates were centrifuged for 3 min at 17,000 g and filtered with 0,8 μm PES filters (Sartorius). Filtrates were treated with benzonase (Novagen) and micrococcal nuclease (New England Biolabs) at 37 °C for 2 h to remove the free-floating nucleic acids. Subsequently, samples were extracted using the QIAamp Viral RNA Mini Kit (Qiagen) according to the manufacturer’s instructions, without addition of carrier RNA to the lysis buffer. Reverse transcription and second strand synthesis was performed by an adjusted version of the Whole Transcriptome Amplification (WTA2) protocol as described previously (Sigma-Aldrich) (68). Sequencing library was constructed with the Nextera XT Library Preparation Kit (Illumina). The size of the library was checked with Bioanalyzer (Agilent Technologies) with a High Sensitivity DNA chip and the 2nM pooled libraries were sequenced on an Illumina NextSeq 500 platform (2×150bp paired-end).

### Data analysis

Low quality reads, ambiguous bases, primer and adapter sequences were removed from the paired-end reads with Trimmomatic v0.36 with default parameters (69). Trimmed reads were *de novo* assembled with metaSPAdes from SPAdes software v3.11.1 using 21, 33, 55, 77 k-mer lengths (70). The obtained contigs were annotated with DIAMOND v0.9.10 against a non-redundant protein database (71). The contigs annotated as “Rotavirus” were further investigated using the nucleotide BLAST against a nucleotide reference database to identify the gene segments (72). The incomplete contigs were completed *in silico* by mapping the trimmed reads of corresponding samples against the reference sequence determined by the highest BLASTn nucleotide similarity with the lowest e-value using BWA software v0.5.9 (73) and SAMtools v1.6 (74). Open reading frames were determined by the web-based NCBI ORF Finder tool (75) (www.ncbi.nlm.nih.gov/orffinder).

### Assignment of GCs and phylogenetic analyses

The genotypes were assigned using RotaC tool (http://rotac.regatools.be). The sequences whose genotypes could not be determined were sent to the RCWG for assignment of novel genotypes.

Reference strains were downloaded from Genbank in order to represent all the relevant genotypes per gene segment. Codon-based nucleotide level multiple sequence alignments were done using MUSCLE (76) with default parameters in MEGA software v7.0.26 (77). Pairwise nucleotide distances were calculated using number of identical residues in relation to the length of the alignment with bio3d package in R (78). Alignments were trimmed with trimAL v1.2 with automated1 parameter (79). Optimized number of bootstrap replicates (100 to 1000) were determined by the autoMRE option and maximum likelihood trees were reconstructed with RaxML-NG (80). GTR+G+I nucleotide substitution model is used for trees of all segments, except for NSP4 and NSP5, as they did not converge after 1000 bootstraps under the GTR+G+I model (TIM3+I+G and HKY+I+G, respectively). FigTree v1.4.3 from the BEAST package was used for phylogenetic tree visualization and manipulation (81). The GCs were illustrated on a world map using the maps package in R software (82).

### Data availability

The data have been deposited with links to BioProject accession number PRJNA562472 in the NCBI BioProject database (https://www.ncbi.nlm.nih.gov/bioproject/). The data is also deposited to GenBank under the following accession numbers: MN433617-27 (BB89-15), MN539284-94 (BR89-60), MN528116-26 (GKS-897), MN477236-46 (GKS-912), MN528101-15 (GKS-926), MN528075-85 (GKS-929), MN528086-MN528100 (GKS-934), MN551587-97 (GKS-941), MN477225-35 (GKS-954), MN551598-MN551608 (KCR10-93), MN567261-72 (K212). Reference strains that were used to construct the multiple sequence alignments are listed in Supplementary Table S5.

### Ethical Statement

Bat capture and sampling were conducted with the permissions of the Wildlife and Hunting Department of the Gabonese Ministry of Water and Forestry (N°003/MEFE-PA/ SG/DGEF/DCF and N°0021/MEFE-PA/SG/DGEF/DCF), and under clearance 314/5327.74.1.6 from the State Office of Energy and Agriculture, the Environment and Rural Areas Schleswig-Holstein (LANU) and clearances 133/24.03.2008 and 192/ 26.03.2009 from the Bulgarian Ministry of Environment and Water. For the Ghanaian bats, ethics approval was obtained from the Committee for Human Research, Publications and Ethics of Komfo Anokye Teaching Hospital and School of Medical Sciences, Kwame Nkrumah University of Science and Technology, Kumasi. Research samples were exported under a state agreement between the Republic of Ghana and the Federal Republic of Germany, represented by the City of Hamburg. Additional export permission was obtained from the Veterinary Services of the Ghana Ministry of Food and Agriculture.

## Acknowledgements

This project has received funding from the European Union’s Horizon 2020 research and innovation program under the Marie Skłodowska-Curie agreement No 721367 granted to J.F.D, J.M and M.V.R. EU Horizon 2020 projects EVAg (grant agreement number 653316) and COMPARE (agreement number 643476) granted to C.D, and the Russian Science Foundation grant 19-15-00055 to A.N.L also provided funding to the current study. H.U.E. had a personal scholarship from the BONFOR intramural program at the University of Bonn. German Federal Ministry of Education and Research (BMBF) (project code 01KIO16D), Deutsche Forschungsgemeinschaft (DFG DR 772-3/1), Deutsche Forschungsgemeinschaft within the Africa Infectious Diseases program through grants to C.D and Y.A.S (DR 772/3-1) and to S.O (KA1241/18-1) were also among the funding contributions. A personal scholarship granted to A. R from the German Academic Exchange Service (DAAD) supported field work in Costa Rica. D.J is supported by the ‘Fonds Wetenschappelijk Onderzoek’ (Research foundation Flanders (1S78019N). L.B was supported by the ‘Fonds Wetenschappelijk Onderzoek’ (1S61618N). K.C.Y was funded by the Interfaculty Council for Development Cooperation (IRO) from the KU Leuven. The computing power in this work was provided by the VSC (Flemish Supercomputer Centre), financed by the FWO and the Flemish government – department EWI.

## Supplementary Material Legends

**Table S1.** RT-PCR oligonucleotides for the initial rotavirus screening against VP1

**Table S2.** Taxonomical annotation, sampling time and location, RVA PCR detection information of the bat samples

**Table S3.** RVA-positive bat samples detected by targeted RT-PCR and undergone viral metagenomics

**Table S4.** Examples of reassortments and unusual genotype constellations among bat RVA strains and distinct RVA genotype constellations in the same bat species

**Table S5.** The Genbank accession numbers of the reference RVA strains used in the study

**Figure S1.** RVA-positive bat families and species

**Figure S2.** Heatmap of pairwise nucleotide identities (NI) of the unusual RVA strains

Supplementary Material and Methods

