## Supplemental data for "At least seven distinct rotavirus genotype constellations in bats with evidence of reassortment and zoonotic transmissions"

**Table S1.** RT-PCR oligonucleotides for the initial rotavirus screening against VP1

| ID no. | Sequence (5' → 3') | Position | Genome segment | Polarity | Assay type |
|---|---|---|---|---|---|
| PanRota-F1570 | TAYACIGAYGTITCICARTGGGA | 1570-1593[a] | VP1 | + | Heminested RT-PCR, 1st round |
| PanRota-R1922 | GCGTAGTTGTCGTCICCRTCBAC | 1900-1922[a] | VP1 | - | 1st and 2nd rd |
| PanRota-F1585a | CARTGGGATTCGTCICAGCAYAAYAC | 1585-1610[a] | VP1 | + | 2nd rd |
| PanRota-F1585b | CARTGGGACGCCAGICAACATAAYAC | 1585-1610[a] | VP1 | + | 2nd rd |

ID, identification; RT-PCR, reverse transcription–PCR; [a]corresponding to Rotavirus A G11P[25] Dhaka6 VP1 (GenBank # EF560705); Variant forms of primers (marked consecutively with an alphabetic character in the last position) were mixed together equally and from then on treated as one single primer.

**Table S2.** Taxonomical annotation, sampling time and location, RVA PCR detection information of the bat samples

| Order-Family | Species | No. of samples per sampling site and year | | | | | | | | PCR positive (%) | Positive samples (ID) |
|---|---|---|---|---|---|---|---|---|---|---|---|
| | | **total** | **BGR 2008** | **BGR 2009** | **CRC 2010** | **GAB 2009** | **DEU 2008** | **GHA 2009** | **ROU 2008** | | |
| **Chiroptera-Pteropodidae** | *Eidolon helvum* | **226** | | | | | | 226 | | 1 (0.4%) | K212 |
| | *Micropteropus pusillus* | **1** | | | | | | 1 | | 0 (0%) | |
| | *Rousettus aegyptiacus* | **10** | | | | 8 | | 2 | | 0 (0%) | |
| **Chiroptera-Rhinolophidae** | *Rhinolophus blasii* | **90** | 82 | 8 | | | | | | 1 (1.1%) | BB89-15 |
| | *Rhinolophus euryale* | **336** | 244 | 92 | | | | | | 2 (0.6%) | BBR89-2, BR89-60 |
| | *Rhinolophus ferrum-equinum* | **52** | 45 | 6 | | | | | 1 | 0 (0%) | |
| | *Rhinolophus hipposideros* | **6** | 6 | | | | | | | | |
| | *Rhinolophus landeri* | **1** | | | | | | 1 | | 0 (0%) | |
| | *Rhinolophus mehelyi* | **22** | 14 | 8 | | | | | | 0 (0%) | |
| | *Rhinolophus spec.* | **6** | | | | 6 | | | | 0 (0%) | |
| **Chiroptera-Hipposideridae** | *Hipposideros cf ruber/caffer* | **183** | | | | 46 | | 137 | | 2 (1.1%) | GKS-637, GKS-660 |
| | *Hipposideros cf spec* | **2** | | | | | | 2 | | 0 (0%) | |
| | *Hipposideros gigas* | **67** | | | | 67 | | | | 10 (14.9%) | GKS-897, GKS-912, GKS-926, GKS-929, GKS-934, GKS-941, GKS-942, GKS-953, GKS-954, GKS-955 |
| | *Hipposideros abae* | **62** | | | | | | 62 | | 0 (0%) | |
| **Chiroptera-Nycteridae** | *Nycteris spec.* | **3** | | | | | | 3 | | 0 (0%) | |
| **Chiroptera-Emballonuridae** | *Coleura afra* | **5** | | | | | | 5 | | 0 (0%) | |
| | *Peropteryx kappleri* | **5** | | | 5 | | | | | 0 (0%) | |
| **Chiroptera-Phyllostomidae** | *Anoura geoffroyi* | **100** | | | 100 | | | | | 0 (0%) | |
| | *Carollia castanea* | **1** | | | 1 | | | | | 0 (0%) | |
| | *Carollia perspicillata* | **203** | | | 203 | | | | | 1 (0.5 %) | KCR10-93 |
| | *Enchisthenes hartii* | **3** | | | 3 | | | | | 0 (0%) | |
| | *Glossophaga commissarisi* | **3** | | | 3 | | | | | 0 (0%) | |
| | *Glossophaga soricina* | **22** | | | 22 | | | | | 0 (0%) | |
| **Chiroptera-Mormoopidae** | *Pteronotus parnellii* | **21** | | | 21 | | | | | 0 (0%) | |
| **Chiroptera-Natalidae** | *Natalus lanatus* | **3** | | | 3 | | | | | 0 (0%) | |
| **Chiroptera-Vespertilionidae** | *Barbastella barbastellus* | **13** | 12 | | | | | | 1 | 0 (0%) | |
| | *Miniopterus inflatus* | **2** | | | | 2 | | | | 0 (0%) | |
| | *Miniopterus schreibersii* | **77** | 39 | | | | | | 38 | 0 (0%) | |
| | *Myotis brandtii* | **17** | | | | | 17 | | | 0 (0%) | |
| | *Myotis alcathoe* | **2** | 2 | | | | | | | 0 (0%) | |
| | *Myotis bechsteinii* | **57** | 32 | | | | 25 | | | 0 (0%) | |
| | *Myotis capaccini* | **1** | 1 | | | | | | | 0 (0%) | |
| | *Myotis dasycneme* | **149** | | | | | 149 | | | 0 (0%) | |
| | *Myotis daubentonii* | **110** | 7 | | | | 103 | | | 1 (0.9%) | SW78-39 |
| | *Myotis emarginatus* | **5** | 5 | | | | | | | 0 (0%) | |
| | *Myotis myotis* | **77** | 3 | | | | 60 | | 14 | 0 (0%) | |
| | *Myotis mystacinus* | **51** | | | | | 51 | | | 0 (0%) | |
| | *Myotis nattereri* | **27** | 2 | | | | 25 | | | 0 (0%) | |
| | *Myotis oxygnathus* | **22** | 1 | | | | | | 21 | 0 (0%) | |
| | *Nyctalus leisleri* | **3** | 3 | | | | | | | 0 (0%) | |
| | *Nyctalus noctula* | **11** | | | | | 2 | | 9 | 0 (0%) | |
| | *Pipistrellus cf nanus/nanulus* | **3** | | | | | | 3 | | 0 (0%) | |
| | *Pipistrellus nathusii* | **2** | | | | | 2 | | | 0 (0%) | |
| | *Pipistrellus pipistrellus* | **37** | | | | | 37 | | | 0 (0%) | |
| | *Pipistrellus pygmaeus* | **29** | 2 | | | | 27 | | | 0 (0%) | |
| | *Pipistrellus spec.* | **6** | | | | | | 6 | | 0 (0%) | |
| | *Plecotus auritus* | **5** | 2 | | | | 3 | | | 0 (0%) | |
| | *Plecotus austriacus* | **1** | | | | | 1 | | | 0 (0%) | |
| **Chiroptera-Molossidae** | *Mops spec.* | **2** | | 1 | | | | 1 | | 0 (0%) | |
| | **Total (46 species)** | **2142** | **502** | **115** | **361** | **129** | **502** | **449** | **84** | **18 (0.8%)** | |

Country: BGR = Bulgaria; CRC = Costa Rica; GAB = Gabon; DEU = Germany; GHA = Ghana; ROU = Romania

**Table S3.** RVA-positive bat samples detected by targeted RT-PCR and undergone viral metagenomics

| Sample | Host | Country | Place | Year |
|---|---|---|---|---|
| BBR89-2 | *Rhinolophus euryale* | Bulgaria | Bratanova | 2008 |
| BB89-15 | *Rhinolophus blasii* | | Elenas Cave | |
| BR89-60 | *Rhinolophus euryale* | | Roman Horse Cave | |
| SW78-39 | *Myotis daubentonii* | Germany | Wahlstorf, SH | 2008 |
| GKS-660 | *Hipposideros caffer* | Gabon | Zadie | 2009 |
| GKS-637 | | | | |
| GKS-897 | *Hipposideros gigas* | Gabon | Faucon | 2009 |
| GKS-912 | | | | |
| GKS-926 | | | | |
| GKS-929 | | | | |
| GKS-934 | | | | |
| GKS-941 | | | | |
| GKS-942 | | | | |
| GKS-953 | | | | |
| GKS-954 | | | | |
| GKS-955 | | | | |
| K212 | *Eidolon helvum* | Ghana | Kumasi | 2009 |
| KCR10-93 | *Carollia perspicillata* | Costa Rica | Orosi | 2010 |

**Table S4**. a. Examples of reassortments among bat RVA strains, b. Examples of bat RVA strains with unusual genotype constellations, potentially resulting from (multiple) reassortment events, c. Examples of distinct RVA genotype constellations in the same bat species

| a. Strains | VP7 | VP4 | VP6 | VP1 | VP2 | VP3 | NSP1 | NSP2 | NSP3 | NSP4 | NSP5 | Host Species | Host Family | Diet |
|---|---|---|---|---|---|---|---|---|---|---|---|---|---|---|
| **RVA/Bat-wt/BGR/BB89-15/2008/G3P[3]** | G3 | P[3] | I3 | R3 | C3 | M3 | A9 | N3 | T3 | E3 | H6 | *Rhinolophus blasii* | *Rhinolophidae* | I |
| **RVA/Bat-wt/BGR/BB89-60/2008/G3P[3]** | G3 | P[3] | I3 | R3 | C3 | M3 | A9 | N3 | T3 | E3 | H6 | *Rhinolophus euryale* | *Rhinolophidae* | I |
| RVA/Bat-wt/CHN/LZHP2/2015/G3P[3] | G3 | P[3] | I3 | R3 | C3 | M3 | A9 | N3 | T3 | E3 | H6 | *Hipposideros pomona* | *Hipposideridae* | I |
| RVA/Bat-tc/CHN/MSLH14/2012/G3P[3] | G3 | P[3] | I8 | R3 | C3 | M3 | A9 | N3 | T3 | E3 | H6 | *Rhinolophus hipposideros* | *Rhinolophidae* | I |
| RVA/Bat-tc/CHN/MYAS33/2013/G3P[10] | G3 | P[10] | I8 | R3 | C3 | M3 | A9 | N3 | T3 | E3 | H6 | *Aselliscus stoliczkanus* | *Hipposideridae* | I |
| RVA/Bat-wt/CHN/BSTM70/2015/G3P[3] | G3 | P[3] | I8 | R3 | C3 | M3 | A29 | N3 | T3 | E3 | H6 | *Taphozous melanopogon* | *Emballonuridae* | I/F |
| RVA/Bat-wt/CHN/YSSK5/2015/G3P[3] | G3 | P[3] | I8 | R20 | C2 | M1 | A9 | N3 | T3 | E3 | H6 | *Scotophilus kuhlii* | *Vespertilionidae* | I |

| b. Strains | VP7 | VP4 | VP6 | VP1 | VP2 | VP3 | NSP1 | NSP2 | NSP3 | NSP4 | NSP5 | Host Species | Host Family | Diet |
|---|---|---|---|---|---|---|---|---|---|---|---|---|---|---|
| RVA/Bat-wt/ZMB/LUS12-14/2012/G3P[3] | G3 | P[3] | I3 | R2 | C2 | M3 | A9 | N2 | T3 | E2 | H3 | *Rhinolophus simulator* | Rhinolophidae | I |
| RVA/Bat-wt/CHN/YSSK5/2015/G3P[3] | G3 | P[3] | I8 | R20 | C2 | M1 | A9 | N3 | T3 | E3 | H6 | *Scotophilus kuhlii* | Vespertilionidae | I |
| RVA/Bat/KEN/322/Kwale/2015/G3P[10] | G3 | P[10] | I2 | R8 | C3 | M5 | A5 | N3 | T6 | E3 | H6 | *Taphozous mauritianus* | Emballonuridae | I |

| c. Strains | VP7 | VP4 | VP6 | VP1 | VP2 | VP3 | NSP1 | NSP2 | NSP3 | NSP4 | NSP5 | Host Species | Host Family | Diet |
|---|---|---|---|---|---|---|---|---|---|---|---|---|---|---|
| RVA/Bat-wt/CMR/BatLy17/2014/G30P[47] | G30 | P[47] | I22 | R15 | C15 | M14 | A25 | N15 | T17 | E22 | H17 | *Eidolon helvum* | *Pteropodidae* | F |
| **RVA/Bat-wt/GHA/K212/2009/G30P[47]** | G30 | P[47] | I22 | R15 | C15 | M14 | A25 | N15 | T17 | E22 | H17 | *Eidolon helvum* | *Pteropodidae* | F |
| RVA/Bat-wt/CMR/BatLy03/2014/G25P[43] | G25 | P[43] | I15 | R16 | C8 | M15 | A26 | N8 | T11 | E23 | H10 | *Eidolon helvum* | *Pteropodidae* | F |
| RVA/Bat/SAU/KSA402/2012/G25P[43] | G25 | P[43] | I15 | R16 | C8 | M15 | A26 | N8 | T11 | E23 | H10 | *Eidolon helvum* | *Pteropodidae* | F |

### **Table S5**. The Genbank accession numbers of the reference RVA strains used in the study

| Strains | VP1 | VP2 | VP3 | VP4 | NSP1 | VP6 | NSP3 | VP7 | NSP2 | NSP4 | NSP5 |
|---|---|---|---|---|---|---|---|---|---|---|---|
| RVA/Alpaca-tc/PER/SA44/2014/G3P40 | | | | KT935478 | | | | | | | |
| RVA/Alpaca-wt/PER/356/2010/G3P14 | | | | KT878993.1 | | | | | | | |
| RVA/Alpaca-wt/PER/Alp11B/2010/G35P50 | | | | KY971955.1 | | | | KY971977.1 | | | |
| RVA/Bat-wt/KEN/322/Kwale/2015/G3P10 | MH285826.1 | MH285827.1 | MH285828.1 | MH285829.1 | MH285830.1 | MH285831.1 | MH285832.1 | MH285834.1 | MH285833.1 | MH285835.1 | MH285836.1 |
| RVA/Bat-wt/KEN/BATp39/2015/G36P51 | MH285837.1 | MH285838.1 | MH285839.1 | MH285840.1 | MH285841.1 | MH285842.1 | MH285843.1 | MH285845.1 | MH285844.1 | MH285846.1 | MH285847.1 |
| RVA/Bat-wt/SAU/KSA402/2012/G25P43 | KX420939.1 | KX420940.1 | KX420941.1 | KX420942.1 | KX420943.1 | KX420944.1 | KX420947.1 | KX420946.1 | KX420945.1 | KX420949.1 | KX420948.1 |
| RVA/Bat-tc/CHN/MSLH14/2012/G3P3 | KC960619.1 | KC960620.1 | KC960621.1 | KC960622.1 | KC960625.1 | KC960623.1 | KC960627.1 | KC960626.1 | KC960624.1 | KC960628.1 | KC960629.1 |
| RVA/Bat-tc/CHN/MYAS33/2013/G3P10 | KJ020891.1 | KJ020892.1 | KJ020893.1 | KF649187.1 | KJ020887.1 | KJ020894.1 | KJ020889.1 | KF649188.1 | KJ020888.1 | KJ020890.1 | KF649186.1 |
| RVA/Bat-wt/BRA/3081/2013/G20Px | | | | KR106166.1 | | | KR106164.1 | KR106163.1 | | | KR106165.1 |
| RVA/Bat-wt/BRA/4754/2013/G3P3 | | | | KR106161.1 | | | | KR106162.1 | | KR106159.1 | KR106160.1 |
| RVA/Bat-wt/CHN/BSTM70/2015/G3P3 | KX814924.1 | KX814925.1 | KX814926.1 | KX814922.1 | KX814927.1 | KX814923.1 | KX814928.1 | KX814929.1 | KX814921.1 | KX814930.1 | KX814931.1 |
| RVA/Bat-wt/CHN/GLRL1/2005/G33P48 | KX814935.1 | KX814936.1 | KX814937.1 | KX814933.1 | | KX814934.1 | KX814939.1 | KX814932.1 | KX814938.1 | KX814941.1 | KX814940.1 |
| RVA/Bat-wt/CHN/LZHP2/2015/G3P3 | KX814945.1 | KX814946.1 | KX814947.1 | KX814943.1 | KX814948.1 | KX814944.1 | KX814950.1 | KX814942.1 | KX814949.1 | KX814951.1 | KX814952.1 |
| RVA/Bat-wt/CHN/YSSK5/2015/G3P3 | KX814956.1 | KX814957.1 | KX814958.1 | KX814954.1 | KX814959.1 | KX814955.1 | KX814961.1 | KX814953.1 | KX814960.1 | KX814962.1 | KX814963.1 |
| RVA/Bat-wt/CMR/BatLi08/2014/G31P42 | KX268765.1 | KX268766.1 | KX268767.1 | KX268768.1 | KX268771.1 | KX268769.1 | KX268773.1 | KX268770.1 | KX268772.1 | KX268774.1 | KX268775.1 |
| RVA/Bat-wt/CMR/BatLi09/2014/G30P42 | KX268754.1 | KX268755.1 | KX268756.1 | KX268757.1 | KX268760.1 | KX268758.1 | KX268762.1 | KX268759.1 | KX268761.1 | KX268763.1 | KX268764.1 |
| RVA/Bat-wt/CMR/BatLi10/2014/G30P42 | KX268743.1 | KX268744.1 | KX268745.1 | KX268746.1 | KX268749.1 | KX268747.1 | KX268751.1 | KX268748.1 | KX268750.1 | KX268752.1 | KX268753.1 |
| RVA/Bat-wt/CMR/BatLy03/2014/G25P43 | KX268776.1 | KX268777.1 | KX268778.1 | KX268779.1 | KX268782.1 | KX268780.1 | KX268784.1 | KX268781.1 | KX268783.1 | KX268785.1 | KX268786.2 |
| RVA/Bat-wt/CMR/BatLy17/2014/G30P47 | KX268787.1 | KX268789.1 | KX268790.1 | KX268788.1 | KX268793.1 | KX268791.1 | KX268795.1 | KX268792.1 | KX268794.1 | KX268796.1 | KX268797.1 |
| RVA/Bat-wt/KEN/KE4852/2007/G25P6 | | GU983673.1 | | GU983674.1 | | GU983675.1 | GU983678.1 | GU983676.1 | GU983677.1 | GU983679.1 | GU983680.1 |
| RVA/Bat-wt/ZMB/LUS12-14/2012/G3P3 | LC158119.1 | LC158120.1 | LC158121.1 | LC158117.1 | LC158122.1 | LC158118.1 | LC158116.1 | LC158123.1 | LC158124.1 | LC158125.1 | LC158126.1 |
| RVA/Bat-wt/ZMB/ZFB14-126/2014/GxPx | | | | | | LC277165.1 | LC277163.1 | | LC277162.1 | LC277164.1 | |
| RVA/Bat-wt/ZMB/ZFB14-135/2014/G31Px | LC277168.1 | | | | | LC277169.1 | LC277167.1 | LC277170.1 | | | |
| RVA/Bat-wt/ZMB/ZFB14-52/2014/G31Px | | | | | | LC277160.1 | LC277159.1 | LC277161.1 | | | |
| RVA/Camel/KUW/s21/2010/G10P15 | | | | JX968470.2 | | | | | | | |
| RVA/Camel-wt/SDN/MRC-DPRU447/2009/G8P1 | | | | | KC257086.1 | | | | | KC257089.1 | |
| RVA/Chicken-tc/DEU/02V0002G3/2002/G19P30 | FJ169853.1 | FJ169854.1 | FJ169855.1 | FJ169856.1 | FJ169857.1 | FJ169858.1 | FJ169859.1 | FJ169861.1 | FJ169860.1 | FJ169862.1 | FJ169863.1 |
| RVA/Chicken-tc/GBR/Ch-1/197x/G19P17 | | | | | | D82970.1 | | AB080738.1 | | | |
| RVA/CommonGull-wt/JPN/Ho374/2013/G28P39 | LC088218.1 | LC088219.1 | LC088220.1 | LC088221.1 | LC088224.1 | LC088222.1 | LC088225.1 | LC088226.1 | LC088223.1 | LC088227.1 | LC088228.1 |
| RVA/Cow-tc/GBR/PP-1/1976/G3P7 | | | | | | | | AF427124.1 | | AF427521.1 | |
| RVA/Bovine-tc/USA/UK/1984/G6P5 | | | | JF693051.1 | | | | | | | |
| RVA/Cow-tc/IND/Hg18/1995/G15P21 | | | | AF237665.1 | | | | AF237666.1 | | | |
| RVA/Cow-tc/JPN/Dai-10/2007/G24P33 | AB573070.1 | AB573071.1 | AB573072.1 | AB513836.1 | AB573074.1 | AB573073.1 | AB573075.1 | AB573076.1 | AB513837.1 | AB573077.1 | AB573078.1 |
| RVA/Cow-tc/THA/A5-13/G8P1 | | | | LC133528.1 | | | | | | | |
| RVA/Cow-tc/USA/B223/G10P11 | | | | LC133550.1 | | | | | | | |
| RVA/Cow-tc/USA/NCDV/1971/G6P1 | DQ870493.1 | DQ870494.1 | | | | | | | | | |
| RVA/Cow-tc/USA/WC3/1981/G6P5 | EF560615.1 | EF560616.1 | EF560617.1 | | EF990699.1 | | EF990701.1 | | EF990700.1 | | EF990702.1 |
| RVA/Cow-wt/JPN/Azuk-1/2006/G21P29 | | | | LC553631.1 | | | LC553636.1 | | | | |

| | | | | | | | | | | | |
|---|---|---|---|---|---|---|---|---|---|---|---|
| RVA/Dog-wt/HUN/135/2012/G3P3 | KJ875791.1 | KJ875792.1 | KJ875793.1 | KJ875794.1 | KJ875797.1 | KJ875795.1 | KJ875799.1 | KJ875796.1 | KJ875798.1 | KJ875800.1 | KJ875801.1 |
| RVA/Dog-tc/ITA/RV198-95/1995/G3P3 | HQ661134.1 | HQ661135.1 | HQ661136.1 | HQ661137.1 | HQ661140.1 | HQ661138.1 | HQ661142.1 | HQ661139.1 | HQ661141.1 | HQ661143.1 | HQ661144.1 |
| RVA/Guanaco-wt/ARG/Chubut/1999/G8P14 | FJ347100.1 | FJ347101.1 | FJ347102.1 | FJ347103.1 | FJ347106.1 | FJ347104.1 | FJ347108.1 | FJ347105.1 | FJ347107.1 | FJ347109.1 | FJ347110.1 |
| RVA/Horse-tc/USA/FI14/1981/G3P12 | | | | | KM454487.1 | | | | | | |
| RVA/Horse-tc/GBR/H-2/1976/G3P12 | | | | KM454495.1 | | | | | | | |
| RVA/Horse-tc/GBR/L338/1991/G13P18 | JF712555.1 | JF712556.1 | JF712557.1 | JF712558.1 | JF712561.1 | JF712559.1 | JF712560.1 | JF712562.1 | JF712563.1 | JF712564.1 | JF712565.1 |
| RVA/Horse-tc/USA/FI23/1981/G14P12 | | | | | | | | KM454508.1 | | | |
| RVA/Horse-wt/ARG/E30/1993/G3P12 | | | | | | | | | | | JF712576.1 |
| RVA/Horse-wt/ARG/E3198/2008/G3P3 | JX036365.1 | JX036366.1 | JX036367.1 | JX036368.1 | JX036371.1 | JX036369.1 | JX036373.1 | JX036370.1 | JX036372.1 | JX036374.1 | JX036375.1 |
| RVA/Human-wt/ITA/ME848/2012/G12P8 | KR632623.1 | KR632624.1 | KR632625.1 | KR632621.1 | KR632626.1 | KR632622.1 | KR632628.1 | KR632620.1 | KR632627.1 | KR632629.1 | KR632630.1 |
| RVA/Human/CHN/ZTR-5/XXXX/G3P2 | JF896465.1 | JF896466.1 | JF896467.1 | JF896468.1 | JF896471.1 | JF896469.1 | JF896473.1 | JF896470.1 | JF896472.1 | JF896474.1 | JF896475.1 |
| RVA/Human-tc/KEN/B10/1987/G3P2 | HM627553.1 | HM627554.1 | HM627555.1 | HM627556.1 | HM627559.1 | HM627557.1 | HM627561.1 | HM627558.1 | HM627560.1 | HM627562.1 | HM627563.1 |
| RVA/Human-tc/CHN/L621/2006/G3P9 | JX946159.1 | JX946160.1 | JX946161.1 | EU708574.1 | JX946163.1 | JX946162.1 | JX946165.1 | EU708588.1 | JX946164.1 | JX946166.1 | JX946167.1 |
| RVA/Human-tc/GBR/A64/1987/G10P1114 | | | | | | | | EF672567.1 | | | |
| RVA/Human-tc/GBR/ST3/1975/G4P2 | | | | | | | | EF672616.1 | | | |
| RVA/Human-tc/IND/116E/1985/G9P11 | | | | FJ361204.1 | | | | | | | |
| RVA/Human-tc/IND/69M/1980/G8P10 | | | | M60600.1 | | | | EF672560.1 | | | |
| RVA/Human-tc/JPN/AU-1/1982/G3P9 | DQ490533.1 | DQ490536.1 | DQ490537.1 | D10970 | D45244 | DQ490538.1 | DQ490535.1 | D86271.1 | DQ490534.1 | | AB008656 |
| RVA/Human-tc/THA/Mc323/1989/G9P19 | | | | D38052.1 | | | | | | | |
| RVA/Human-tc/THA/T152/1998/G12P9 | | | | | | | | | | | DQ146706.1 |
| RVA/Human-tc/USA/DS-1/1976/G2P1B4 | HQ650116.1 | HQ650117.1 | HQ650118.1 | HQ650119.1 | HQ650120.1 | HQ650121.1 | HQ650122.1 | HQ650124.1 | HQ650123.1 | HQ650125.1 | HQ650126.1 |
| RVA/Human-tc/USA/Wa/1974/G1P1A8 | KT694939.1 | KT694940.1 | KT694941.1 | KT694942.1 | KT694945.1 | KT694943.1 | KT694947.1 | KT694944.1 | KT694946.1 | KT694948.1 | KT694949.1 |
| RVA/Human-tc/USA/WI61/1983/G9P1A8 | | | | | | | | LC482504.1 | | | |
| RVA/Human-wt/BEL/B4106/2000/G3P14 | | | | | | | | | | AY740732.1 | |
| RVA/Human-wt/BEL/BEF06018/2014/G29P41 | | | | KU128895.1 | | | | | | | |
| RVA/Human-wt/BGD/Dhaka6/2001/G11P25 | | | | AY773004.2 | | | | | | | |
| RVA/Human-wt/BRA/QUI-35-F5/2010/G3P9 | | | | | KF185099.1 | KF185107.1 | | | | | |
| RVA/Human-wt/CHN/E2451/2011/G3P9 | JX946168.1 | JX946169.1 | JX946170.1 | JX946171.1 | JX946174.1 | JX946172.1 | JX946176.1 | JX946173.1 | JX946175.1 | JX946177.1 | JX946178.1 |
| RVA/Human-wt/ECU/Ecu534/2006/G20P28 | | | | EU805773.1 | | EU805774.2 | | EU805775.1 | | | |
| RVA/Human-wt/HUN/Hun5/1997/G6P14 | | | | | EF554110.1 | | | | | | |
| RVA/Human-wt/NPL/KTM368/2004/G11P25 | | | | | | GU199496.1 | | | | | |
| RVA/Human-wt/SUR/2014735512/2013/G20P28 | KX257410.1 | KX257409.1 | KX257408.1 | KX257407.1 | KX257415.1 | KX257406.1 | KX257414.1 | KX257413.1 | KX257405.1 | KX257412.1 | KX257411.1 |
| RVA/Human-wt/THA/CMH222/2001/G3P3 | | | | DQ288661.1 | | DQ288659.1 | | AY707792.1 | | DQ288660.1 | |
| RVA/Human-wt/US/09US7118/2009/G3P24 | KF541281.1 | KF541282.1 | KF541283.1 | KF541284.1 | KF541287.1 | KF541285.1 | KF541288.1 | KF541289.1 | KF541286.1 | KF541290.1 | KF541291.1 |
| RVA/Alpaca/PER/ALRVA-Kayra/3386/2010/G3Px | | | | | | | | KT250942.1 | | | |
| RVA/Mouse-tc/UK/EHP/1981/G16P20 | | | | U08424.1 | | | | | | | |
| RVA/Mouse-tc/USA/ETD_822/2007/G16P16 | GQ479947.1 | GQ479948.1 | GQ479949.1 | GQ479950.1 | | | GQ479953.1 | | GQ479954.1 | | GQ479957.1 |
| RVA/Mouse-tc/USA/EW/XXXX/G16P16 | | | | U08429.1 | U08428.1 | U36474.1 | | U08430.1 | | U96335.1 | |
| RVA/Pheasant-tc/GER/10V0112H5/2010/G23P37 | | | | JX204814.1 | | | | | | | |
| RVA/Pheasant-wt/HUN/Phea14246/2008/G23Px | | | | | | | | FN393054.1 | | | |

| | | | | | | | | | | | |
|---|---|---|---|---|---|---|---|---|---|---|---|
| RVA/Pigeon-tc/JPN/PO-13/1983/G18P17 | AB009629.2 | AB009630.2 | AB009631.2 | AB009632.2 | AB009633.2 | D16329.2 | AB009626.2 | D82979.2 | AB009625.2 | AB009627.1 | AB009628.1 |
| RVA/Pig-tc/USA/OSU/1975/G5P7 | | | | KR052770.1 | | | | KR052772.1 | | | |
| RVA/Pig-wt/XXX/A46/XXXX/GxP13 | | | | AY050274.1 | | | | | | | |
| RVA/Pig-wt/BGD/2014016006/G9P49 | | | | KY905314.1 | | | | | | | |
| RVA/Pig-wt/CAN/CE-M-06-0003/2006/G2P27 | | | | | | GU183245.1 | | | | | |
| RVA/Pig-wt/CHN/NMTL/2008/G9P23 | | | | JF781161.1 | | | | | | | |
| RVA/Pig-wt/IRL/61-07-ire/2007/G2P32 | | | | FJ492835.1 | | | | | | | |
| RVA/Pig-wt/JPN/FGP51/2009/G4P34 | | | | AB571047.1 | | | | | | | |
| RVA/Pig-wt/JPN/TJ41/2010/G26Px | | | | | | | | AB605258.1 | | | |
| RVA/Pig-wt/THA/CMP034/2000/G2P27 | | | | DQ534016.2 | | | | | | | |
| RVA/Rabbit-wt/ITA/160/01/2001/G3P22 | | | | AF526374.1 | | | | | | | |
| RVA/Raccoon-tc/JPN/Rac-311/2011/G34P17 | | LC208546.1 | LC208547.1 | | LC208551.1 | LC208549.1 | LC208553.1 | LC208550.1 | LC208552.1 | | |
| RVA/Rat/CHN/RA116/2013/G3P45 | | | | KU243630.1 | | | | | | | LC208555.1 |
| RVA/Rat-wt/GER/KS-11-573/2011/G3P3 | KJ879448.1 | KJ879449.1 | KJ879450.1 | | KJ879454.1 | KJ879452.1 | KJ879456.1 | | | KJ879457.1 | KJ879458.1 |
| RVA/Rat-wt/ITA/Rat14/2015/G3P3 | | | | | | | | | KX398370.1 | | |
| RVA/Rhesus-tc/USA/TUCH/2002/G3P24 | EF583010.1 | EF583011.1 | EF583012.1 | FJ816611.1 | FJ816612.1 | EF583013.1 | FJ816613.1 | FJ816615.1 | FJ816614.1 | FJ816616.1 | FJ816617.1 |
| RVA/Sheep-wt/CHN/Lp14/xxxx/G10P15 | | | | L11599.1 | | L11595.1 | | | | | |
| RVA/Shrew-wt/CHN/LW9/2013/G28P46 | KU243542.1 | KU243571.1 | KU243598.1 | KU243621.1 | KU243377.1 | KU243658.1 | KU243457.1 | KU243688.1 | KU243430.1 | KU243476.1 | KU243505.2 |
| RVA/Simian-tc/USA/RRV/1975/G3P3 | EU636924.1 | EU636925.1 | EU636926.1 | EU636927.1 | EU636928.1 | EU636929.1 | EU636930.1 | EU636932.1 | EU636931.1 | EU636933.1 | EU636934.1 |
| RVA/Simian-tc/USA/SA11g4O-5N/G3P1 | | | | EU636934.1 | | | | | | | |
| RVA/Simian-tc/ZAF/SA11-H96/1958/G3P2 | DQ838640.2 | DQ838635.2 | DQ838645.2 | DQ841262.2 | DQ838599.2 | DQ838650.2 | DQ838610.2 | DQ838620.2 | DQ838615.2 | DQ838625.2 | DQ838630.2 |
| RVA/SugarGlider-tc/JPN/SG385/2012/G27P36 | AB971760.1 | AB971761.1 | AB971762.1 | AB971763.1 | AB971766.1 | AB971764.1 | AB971768.1 | AB971765.1 | AB971767.1 | AB971769.1 | AB971770.1 |
| RVA/Turkey-tc/DEU/03V0002E10/2003/G22P35 | | | | JX204825.1 | | | | JX204830.1 | | | |
| RVA/Turkey-tc/IRL/Ty-1/1979/G17P38 | | | | LC088110.1 | | | | S58166.1 | | | |
| RVA/Turkey-tc/IRL/Ty-3/1979/G7P35 | | | | | | | | LC088119.1 | | | |
| RVA/VelvetScoter-tc/JPN/RK1/1989/G18P17 | | | | | LC088102.1 | | | | | | |
| RVA/Vicuna-wt/ARG/C75/2010/G8P14 | | | | | | | | | | JX070055.1 | |

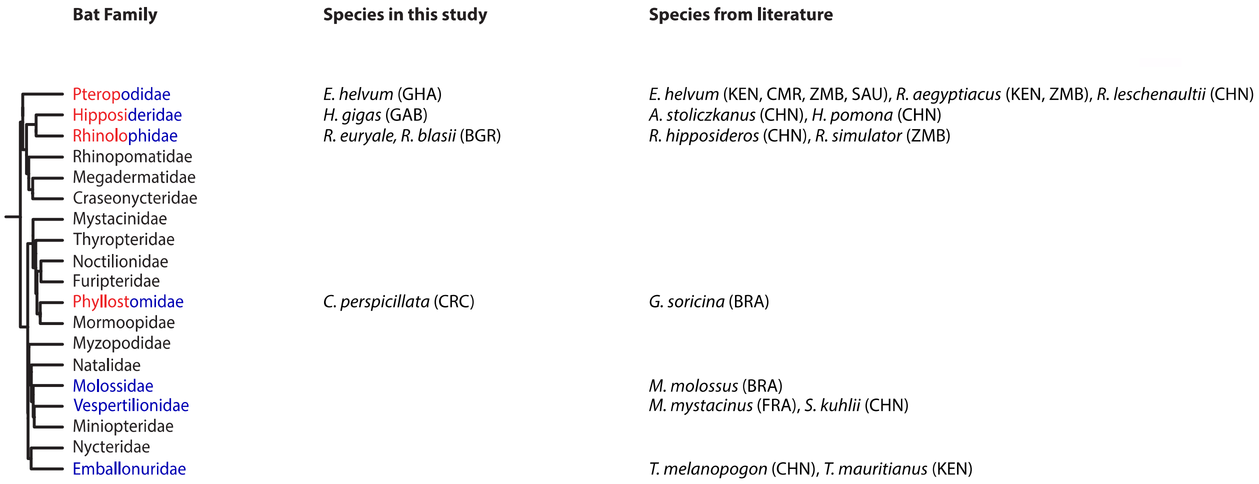


**Figure S1.** RVA-positive bat families and species. The RVA-positive bat families reported in the present study (red) and in literature (blue) are shown on the phylogenetic tree adapted from Simmons et al (2003). No RVA is reported in families in black. A family is accepted positive for the literature group if more than 1 RVA segment was submitted to GenBank. The corresponding bat species and the country of sample collection are also displayed. Country: GHA = Ghana, FRA = France, BRA = Brazil, ZMB = Zambia, SAU = Saudi Arabia, CRC = Costa Rica, KEN = Kenya, CHN = China, BGR = Bulgaria, GAB = Gabon

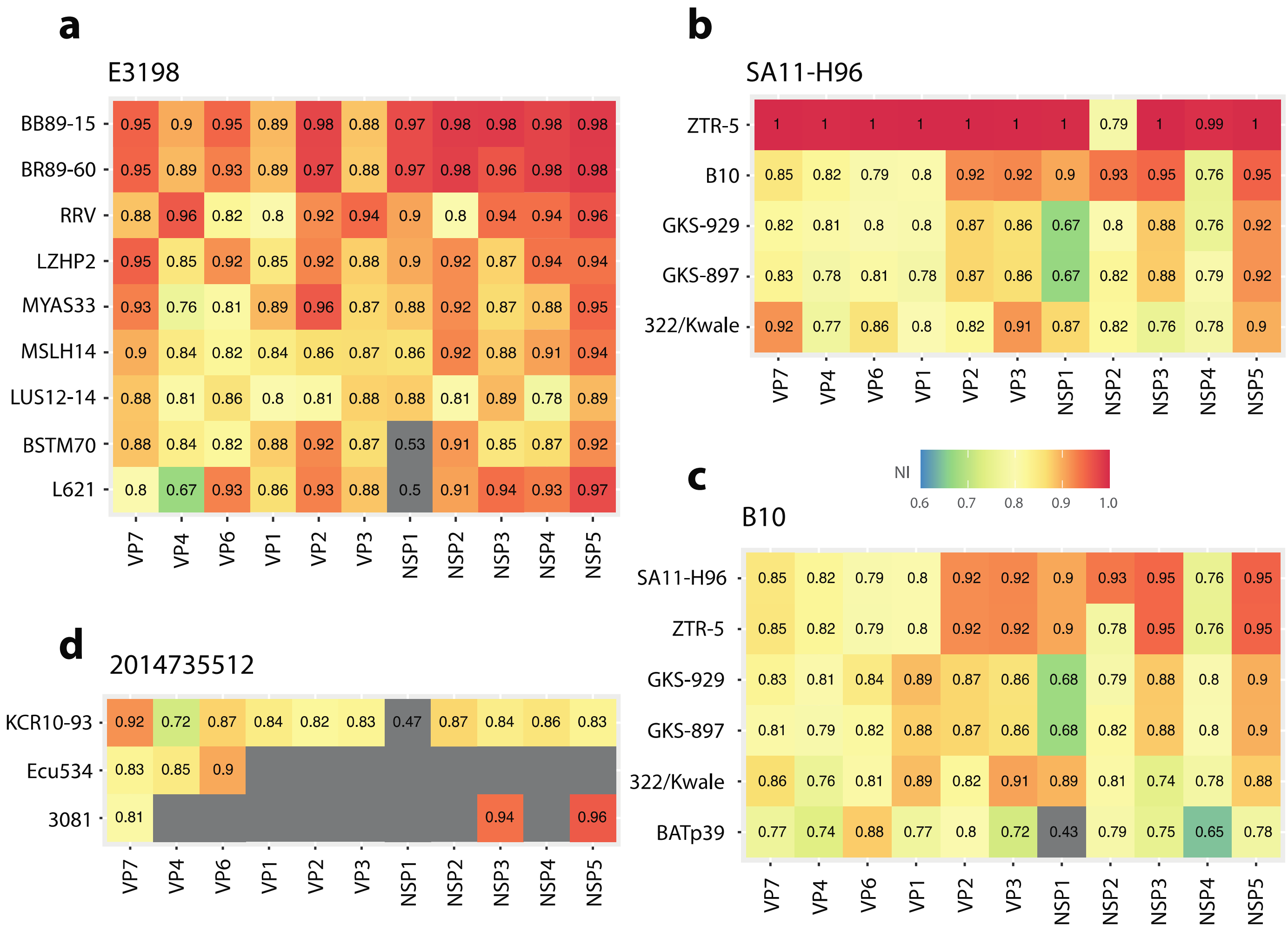


**Figure S2.** Heatmap of pairwise nucleotide identities (NI) of the unusual RVA strains: E3198 (a), SA11-H96 (b), B10 (c), 2014735512 (d). Grey colour indicates the nucleotide identities below 0.6 or lack of sequence information for the compared strain

**Supplementary Material and Methods**

**Screening VP1-Consensus-PCR**

25-µL SuperScript© III with Platinum© Taq DNA Polymerase One-Step RT-PCR reactions as described by the manufacturer (INVITROGEN, Karlsruhe, Germany) used 800 nM each of 1$^{st}$-round primers, 1 µg bovine serum albumin, $MgSO_4$ up to a total concentration of 2.4 mM, plus 5 µL RNA extract. Amplification involved 30 min at 48°C; 3 min at 95°C; 10 cycles of 20 s at 95°C, 20 s starting at 60°C with a decrease of 1°C per cycle, and 35 s at 72°C; 40 cycles of 20 s at 95°C, 20 s at 50°C, and 35 s at 72°C; and a final elongation step of 2 min at 72°C. 50-µL Platinum Taq reactions as described by the manufacturer (INVITROGEN, Karlsruhe, Germany) used 2 µL of 1st-round PCR product, 2 mM $MgCl_2$ and 800 nM of 2$^{nd}$-round forward primer and 400 nM of the reverse primer. Amplification involved 3 min at 95°C; 10 cycles of 15 s at 95°C, 15 s starting at 62°C with a decrease of 1°C per cycle, and 30 s at 72°C; 40 cycles 15s at 95°C, 15 s at 52°C and 30 s at 72°C; and a final elongation step of 2 min at 72°C. All PCR reactions were carried out in an Eppendorf Mastercycler ep gradient S (Eppendorf AG, Hamburg, Germany).
